# A regenerative fibroblast competence field supports multiple cell fates in adult skin regeneration

**DOI:** 10.64898/2026.10.01.756129

**Authors:** Chih-Han Huang, Chih-Kuan Chen, Chen Yuan Kam, Ben Van Handel, Arijita Sarkar, Yen-Chun Ho, Denis Evseenko, Cheng-Ming Chuong, Hans I-Chen Harn

## Abstract

Adult mammalian wounds typically restore tissue integrity through fibrotic repair rather than regeneration of the original architecture. Here, we compared small non-regenerative and large regenerative skin wounds to investigate how fibroblasts acquire regenerative competence and enter alternative cell-fate programs. Integrated single-cell RNA sequencing identified a transient signaling and extracellular-matrix-remodeling niche enriched in large wounds, together with NCAM1-associated competent fibroblast states. Time-resolved analysis positioned these states within a root-like fibroblast manifold connected to hair dermal-condensate, endothelial, pericyte and glial-associated fate-entry programs. Inferred cell-cell communication networks revealed shared matrix and adhesion signals, superimposed with branch-associated inputs, including VEGF signaling toward NCAM1-positive endothelial-transition cells. Lineage tracing showed that PDGFRA- and NCAM1-expressing lineages contributed to vascular- and nerve/glial-associated regenerative compartments. Functionally, partial NCAM1 perturbation reduced vascular organization while increasing hair follicle neogenesis, whereas stronger inhibition impaired both outcomes. These findings identify NCAM1 as a regulator of regenerative competence and fate specification within a regenerative competence field rather than a marker of a single terminal lineage. We propose that regenerative wounds activate an NCAM1-associated fibroblast competence field whose quantitative modulation biases competent fibroblasts toward alternative outcomes.

## Introduction

Adult tissue regeneration is the restoration of the structures, spatial organization and functions lost after injury. This outcome is distinct from repair, which rapidly restores continuity but commonly leaves a fibrotic substitute. Small full-thickness wounds close through contraction, re-epithelialization and matrix deposition, yet the healed tissue lacks the original hair follicles, dermal adipose tissue and normal connective-tissue architecture. By contrast, sufficiently large wounds in mice can enter a regenerative regime in which new hair follicles and adipocytes arise in the wound center (Ito, Yang et al. 2007, Plikus, Guerrero-Juarez et al. 2017) (Guerrero-Juarez, Dedhia et al. 2019). The small-versus-large wound comparison therefore provides more than a difference in injury scale: it exposes a switch from fibrotic repair to regeneration and offers an experimentally accessible system in which to identify the conditions that permit adult tissue to rebuild rather than merely close.

Regeneration cannot be explained by tissue growth alone. Proliferation may replace cell mass, but it does not recreate the diverse cell identities, reciprocal epithelial-mesenchymal interactions or three-dimensional arrangements required for a functional organ. New appendages must acquire dermal condensate and dermal papilla identities; the surrounding tissue must rebuild vessels, mural support and nerve-associated compartments; and these elements must emerge in the correct sequence and location. Regeneration therefore requires a controlled form of cellular reprogramming in which adult cells recover developmental competence, enter transient states and then select context-appropriate fates (Tanaka and Reddien 2011). This does not imply complete dedifferentiation or acquisition of pluripotency. Studies of salamander limbs and mammalian digit tips instead support partial, lineage-biased reprogramming within a permissive regenerative environment (Gerber, Murawala et al. 2018). Defined transcription factors can also redirect fibroblasts toward endothelial-like identities in culture, showing that fibroblast state is experimentally malleable (Margariti, Winkler et al. 2012, Lee, Park et al. 2017). What remains unresolved is how such competence is generated and allocated among alternative fates in an adult wound without forced expression of lineage regulators(Goldman and Poss 2020).

In highly regenerative systems, these decisions occur within a blastema or morphogenetic field rather than in isolated cells(Kragl, Knapp et al. 2009, Lei, Jiang et al. 2023). A blastema is a transient injury-induced assembly of proliferative, responsive cells positioned beneath a specialized wound epithelium (Harn, Davidson et al. 2023). It contains mesenchymal progenitor-like cells together with immune, vascular, neural and epithelial populations (Gay, Kwon et al. 2013, Godwin, Pinto et al. 2013, Nelson, Reddy et al. 2015), and is organized by extracellular matrix remodeling, innervation and short-range and diffusible signals(Kumar, Godwin et al. 2007). Single-cell studies have shown that the blastema is molecularly heterogeneous and that its mesenchymal compartment contains injury-induced states and putative differentiation trajectories rather than one uniformly multipotent population(Gerber, Murawala et al. 2018, Leigh, Dunlap et al. 2018). The regenerative niche can therefore be viewed as a dynamic signaling and material environment that first establishes competence and then channels competent cells into distinct fate-entry programs (Hsu, Li et al. 2014). In skin, competence can be regulated across tissue domains: cyclic dermal BMP–WNT activity establishes competent and refractory regions, while regenerative waves and injury-density-dependent quorum sensing coordinate responses among spatially separated follicles(Plikus, Mayer et al. 2008, Plikus, Baker et al. 2011, Chen, Wang et al. 2015). A central problem is to determine whether adult mammalian wounds assemble an analogous competence field, which cells occupy its root state, and how interactions among its constituents resolve cell fate after reprogramming.

Fibroblasts are spatially diverse and responsive to inflammatory, epithelial, vascular and matrix-derived cues(Driskell, Lichtenberger et al. 2013, Rinkevich, Walmsley et al. 2015). Fibroblasts possess stable positional identities, but regenerative competence may represent a superimposed, injury-induced and reversible state(Abbasi, Sinha et al. 2020) rather than fixed anatomical lineage. During conventional healing they generate and remodel extracellular matrix, contract the wound and support inflammatory resolution, but persistent activation can stabilize myofibroblast states and fibrosis (Rinkevich, Walmsley et al. 2015). During regenerative healing, fibroblast-lineage cells can also contribute to newly formed dermal papillae and adipocytes, and single-cell analyses reveal extensive heterogeneity among wound fibroblasts (Plikus, Guerrero-Juarez et al. 2017, Guerrero-Juarez, Dedhia et al. 2019). These findings suggest that regeneration depends on more than the presence of a particular pre-existing fibroblast subtype. An alternative model is that the wound creates a transient field in which selected fibroblasts acquire regenerative competence before entering fate-biased intermediates. Distinguishing this model from fixed lineage heterogeneity requires temporal state mapping, spatial validation and lineage analysis, because transcriptional proximity alone cannot establish direct conversion.

Tissue mechanics is one of the modulators of the regenerative field (Harn, Ogawa et al. 2019, Harn, Chen et al. 2021, Harn, Wang et al. 2021, Harn, Chiu et al. 2022, Harn, Davidson et al. 2023). Mechanical forces guide cell rearrangement, folding and fate specification during development (Harn, Jiang et al. 2026), and time-varying mechanical and biochemical feedback can drive the self-organization of skin organoids (Lei, Harn et al. 2023) and appendage primordia (Harn, Jiang et al. 2026). In adult wounds, cell contractility, collagen organization, matrix stiffness and tissue tension form feedback loops that reinforce myofibroblast activation and fibrosis (Harn, Ogawa et al. 2019, Harn, Wang et al. 2021, Harn, Chiu et al. 2022). Regenerative systems instead remodel the extracellular matrix to produce spatially heterogeneous mechanical environments. In wound-induced hair follicle neogenesis, the large wound establishes a permissive morphogenetic field, while local symmetry breaking in tissue mechanics accompanies placode and dermal condensate formation; these patterns differ between laboratory and spiny mice (Harn, Wang et al. 2021). Mechanics is therefore likely to define where and when cells can respond to regenerative signals. The molecular interface that translates this tissue-scale context into reversible competence and subsequent fate choice remains less clear.

More broadly, these observations suggest that regenerative competence is controlled not only by the identity of a signal, but also by its magnitude, duration and spatial reach. Injury-density-dependent quorum sensing, selective modulation of gp130 signaling and an intermediate wound-mechanical range each illustrate that regeneration can emerge within a permissive quantitative window rather than through binary pathway activation(Chen, Wang et al. 2015, Harn, Chen et al. 2021, Harn, Chiu et al. 2022, Shkhyan, Flynn et al. 2023). A molecular system capable of integrating these graded biochemical and mechanical inputs could therefore tune both the maintenance of competence and its resolution into alternative fates.

Cell adhesion molecules provide such an interface. Cell-matrix adhesions couple extracellular matrix properties to the cytoskeleton and nucleus, whereas cell-cell adhesions control contact, sorting, collective movement and the duration of receptor-ligand interactions (Harn, Davidson et al. 2023) (Dupont, Morsut et al. 2011). Changes in force can alter adhesion assembly and signaling, while altered adhesion can reorganize multicellular geometry and thereby change the signals that each cell receives. Synthetic adhesion systems are sufficient to program cell sorting and multicellular architecture, demonstrating that adhesion codes can instruct tissue organization rather than simply maintain cohesion (Stevens, Harris et al. 2023). Adhesion-dependent control is also attractive for regenerative reprogramming because it can act transiently and locally: a cell can change its neighbors, mechanical coupling and signaling access without first being driven to pluripotency. This raises the possibility that a native adhesion molecule could function as a quantitative control point, translating graded changes in signaling and mechanical context into different specifications of a competent fibroblast population across regenerative fates.

Neural cell adhesion molecule 1 (NCAM1, also known as CD56) is a strong candidate for this role. NCAM1 is an immunoglobulin-superfamily cell-surface glycoprotein that participates in homophilic and heterophilic adhesion, cell migration and tissue patterning. Alternative isoforms and post-translational polysialylation can shift NCAM1 between relatively adhesive and permissive states(Rutishauser 2008), allowing the same molecular platform to stabilize contacts or facilitate cellular rearrangement. NCAM1 also functions as a signaling co-receptor: its engagement can activate fibroblast growth factor receptors, Fyn, focal adhesion kinase, MAPK and calcium-dependent pathways that regulate survival, motility and differentiation (Ditlevsen, Povlsen et al. 2008). These properties underlie well-established functions in neural development, axon guidance, synaptic plasticity and muscle progenitor differentiation. NCAM is also expressed in regenerating newt limb mesenchyme and wound epidermis, and antibody blockade delays limb regeneration (Maier, Watanabe et al. 1986). Yet whether NCAM1 marks or regulates a regenerative fibroblast state in adult mammalian skin, and whether its quantitative modulation biases the fates that emerge from that state, are unknown.

We therefore hypothesized that large wounds assemble a transient signaling and matrix-remodeling niche that supports an NCAM1-associated fibroblast competence manifold, from which context-dependent fate-entry programs emerge, and that the level or function of NCAM1-mediated adhesion tunes how regenerative output is allocated among vascular and appendage-associated outcomes. This hypothesis leads to five linked questions. How does the cellular and signaling environment of a regenerative large wound differ from that of a fibrotic small wound? Do regenerative fibroblasts converge on a shared root-associated competent state before entering dermal condensate or dermal papilla, endothelial, pericyte and glial-associated programs? Which niche interactions are shared across these trajectories, and which accompany branch-specific fate entry? Do PDGFRA- and NCAM1-lineage cells contribute to these regenerative compartments in vivo? Finally, can partial modulation of NCAM1 shift tissue-level outcomes without uniformly suppressing regeneration?

To address these questions, we compared non-regenerative small wounds with regenerative large wounds and integrated time-resolved single-cell RNA sequencing with spatial molecular validation. We used state, module and trajectory analyses to identify a root-associated competent fibroblast manifold and to map candidate fate-entry programs, while ligand-receptor analysis was used to nominate shared and branch-associated niche interactions. Complementary lineage-tracing experiments tested whether PDGFRA- and NCAM1-lineage cells contribute to multiple regenerative compartments in vivo. Finally, pharmacologic and antibody-based NCAM1 perturbations were used to determine whether changing NCAM1 function alters the balance between vascular organization and hair follicle neogenesis. Together, this strategy tests NCAM1 as a regulator of population-level fate specification within a regenerative wound niche, while separating transcriptional inference from lineage contribution and functional perturbation.

## Results

### Large wounds establish a regenerative fibroblast competence field

To identify cellular features that distinguish regenerative healing from fibrotic repair, we compared full-thickness small wounds, which close without restoring the original appendage-rich architecture, with large wounds capable of wound-induced hair follicle neogenesis (Fig. 1a). Integrated single-cell analysis revealed a marked expansion of the central fibroblast compartment in large wounds together with increased representation of PDGFRA-associated early dermal-condensate, endothelial-, pericyte- and glial-associated populations (Fig. 1b,c and Supplementary Table 1). These comparisons were descriptive because each condition was represented by one sequencing dataset, but they identified a coordinated cellular program associated with the regenerative wound environment rather than expansion of a single terminal population.

**Figure 1.**
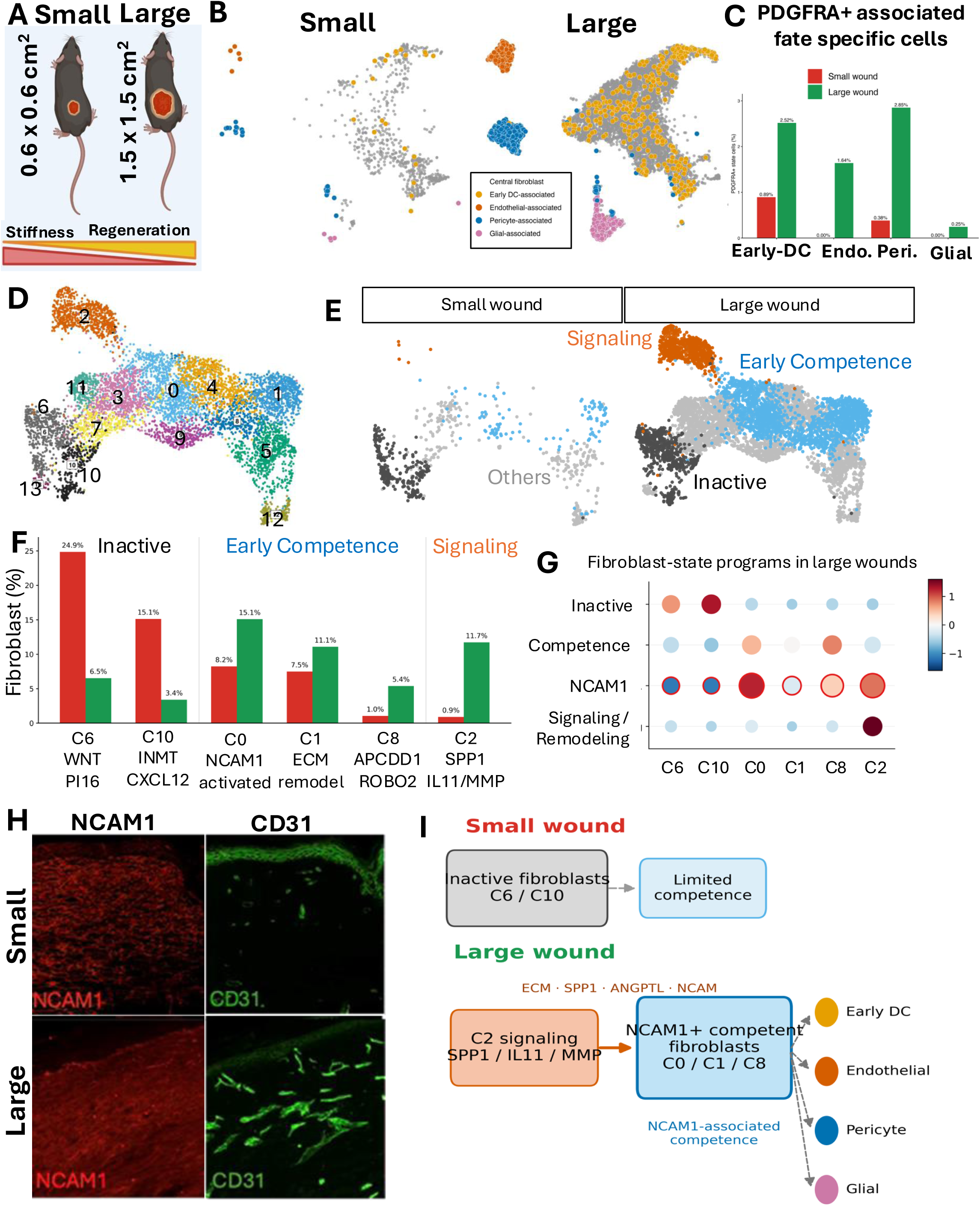
Large wounds contain unique signaling and competent regenerative fibroblasts. a, Schematic of the 0.6 × 0.6 cm small-wound and 1.5 × 1.5 cm large-wound models, which differ in tissue stiffness and regenerative outcome. b, UMAPs of integrated wound cells from small and large wounds, highlighting central fibroblasts and PDGFRA-associated early dermal-condensate (DC), endothelial-, pericyte- and glial-associated populations. Grey denotes other cells. c, Percentages of PDGFRA-associated early-DC, endothelial-, pericyte- and glial-associated cells in small and large wounds. Percentages are descriptive and were calculated within the indicated wound-cell dataset. d, Unsupervised clustering of the integrated fibroblast compartment at resolution 0.9. e, Fibroblast-state landscapes separated by wound condition. Cells were grouped as inactive fibroblasts, early-competence states, signaling-hub fibroblasts or other fibroblast states; the C1 extracellular-matrix (ECM)-remodeling population is indicated. f, Relative abundance of selected fibroblast states within the pooled fibroblast compartment of each wound condition. C6 (WNT2/PI16) and C10 (INMT/CXCL12) represent inactive fibroblasts; C0 is an NCAM1-associated early-competence state; C1 is an ECM-remodeling state; C8 (APCDD1/ROBO2) is associated with competence acquisition; and C2 (SPP1/IL11/MMP-high) is a signaling state. Values above bars show the percentage within each condition. g, Dot plot summarizing inactive/root, competence-acquisition, NCAM1 and signaling/remodeling gene programs across selected large-wound fibroblast clusters. Color denotes mean scaled expression and circle size denotes the fraction of cells expressing the indicated program; NCAM1 is displayed separately and outlined in red. h, Representative immunofluorescence images of NCAM1 and CD31 in small- and large-wound dermis. i, Working model in which large wounds, unlike small wounds, contain a C2 signaling state and an ECM-remodeling niche associated with NCAM1-positive competent fibroblast states (C0/C1/C8) and increased representation of early-DC, endothelial-, pericyte- and glial-associated populations. The single-cell comparisons are population-level associations and do not by themselves establish lineage relationships. The single-cell RNA-sequencing analysis represents one sequencing dataset (n = 1); staining was performed in three biological samples (n = 3).

We next isolated the fibroblast compartment and resolved it into transcriptionally distinct states (Fig. 1d,e and Supplementary Fig. 1a). Two inactive fibroblast populations, C6 (WNT2/PI16) and C10 (INMT/CXCL12), were prominent in small wounds but depleted in large wounds. C6 decreased from 24.9% to 6.5% of wound fibroblasts and C10 decreased from 15.1% to 3.4% (Fig. 1f). Conversely, large wounds were enriched for C0, an NCAM1-associated early-competence state (8.2% versus 15.1%); C1, an extracellular-matrix-remodeling state (7.5% versus 11.1%); C8, a competence-acquisition state marked by APCDD1 and ROBO2 (1.0% versus 5.4%); and C2, a signaling state expressing SPP1, IL11 and matrix metalloproteinases (0.9% versus 11.7%). Independent marker analysis supported these annotations and distinguished the regenerative states from potential contaminating populations (Supplementary Fig. 2 and Supplementary Table 2).

The selected clusters separated along coordinated gene programs rather than by a single marker. C6 and C10 retained inactive or baseline-reticular features, C8 preferentially expressed a competence-acquisition program, and C2 was enriched for signaling and matrix-remodeling genes (Fig. 1g). NCAM1 expression was strongest across competence-associated states, particularly C0, and was more broadly represented in large than small wounds (Fig. 1g and Supplementary Fig. 4a-c). PDGFRA remained detectable across the fibroblast backbone and early fate-associated populations, supporting its use as a broad lineage handle while also showing that PDGFRA transcript expression alone could not establish ancestry (Supplementary Fig. 3). Immunofluorescence confirmed greater NCAM1 abundance in large-wound dermis and showed this field in the context of an expanded CD31-positive vascular network (Fig. 1h and Supplementary Fig. 4d,e). Together, these data indicate that large wounds replace inactive fibroblast-dominated repair with a signaling, matrix-remodeling and NCAM1-associated competence field linked to the emergence of multiple regenerative populations (Fig. 1i).

### A transient signaling and matrix remodeling niche supports fibroblast competence

We asked how the regenerative field developed during the period preceding overt tissue patterning. Integration of fibroblast states from post-wounding days (PWD) 12, 13 and 14 showed that signaling, extracellular-matrix-remodeling and competent states were most prominent at the earlier time points, whereas inactive and other branch-associated states accounted for a greater fraction of the PWD14 landscape (Fig. 2a,b). Alluvial mapping of the day-specific source clusters onto the integrated state assignments showed that several transcriptionally distinct inputs converged on the competent and remodeling compartments (Fig. 2c). Thus, the competence field was not restricted to one static cluster or one obligatory linear precursor. Instead, it formed a transient manifold containing related states with complementary functions.

**Figure 2.**
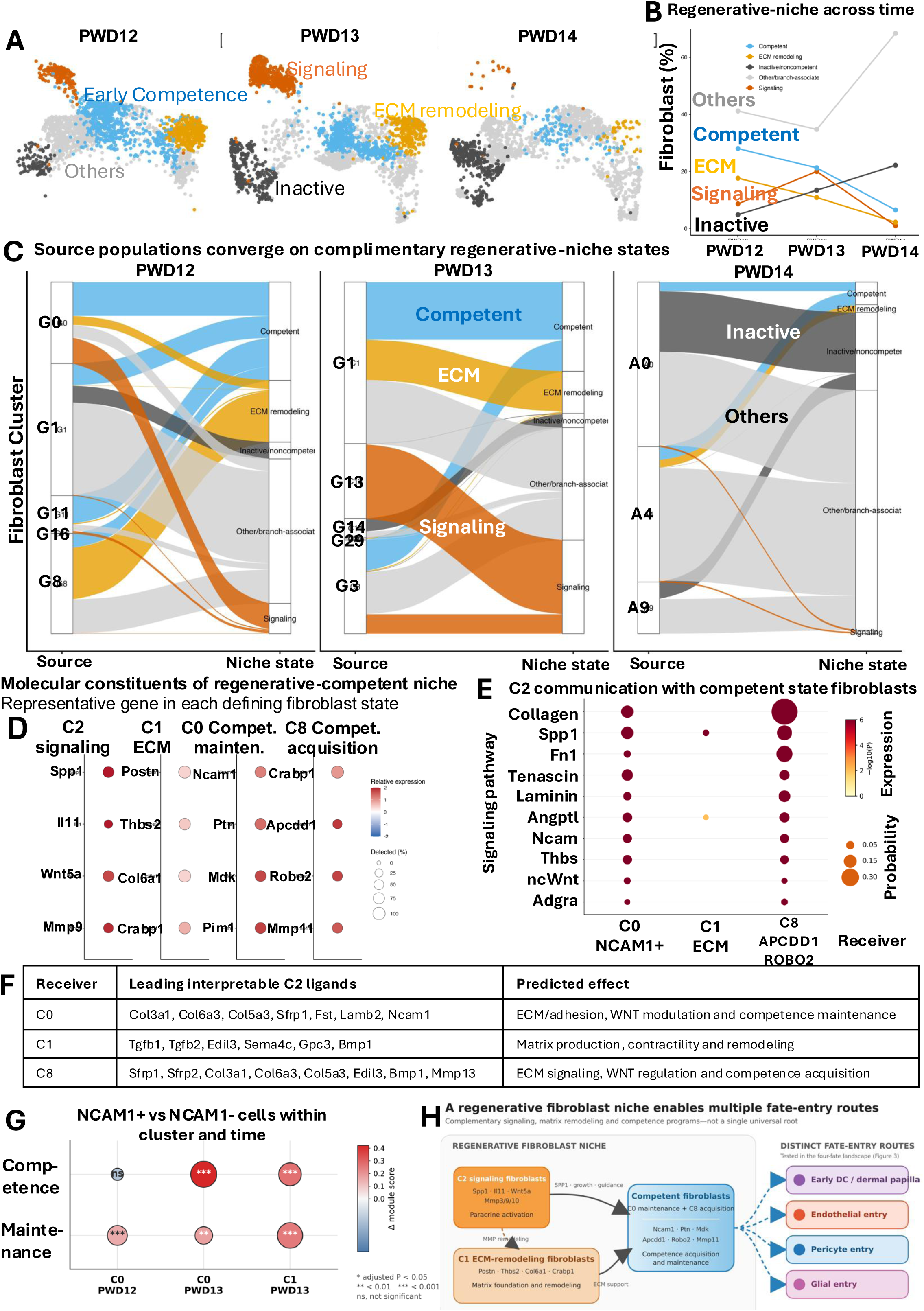
A transient signaling and matrix-remodeling niche supports fibroblast competence. a, UMAPs of the integrated fibroblast-state landscape at post-wounding days (PWD) 12, 13 and 14. Cells are grouped as inactive/noncompetent, signaling, ECM-remodeling, competent or other/branch-associated states. b, Relative composition of these state groups across PWD12–14, expressed as the percentage of fibroblasts within each day. c, Alluvial plots relating source clusters from each day to the integrated regenerative-niche state assignments. Ribbon width represents the number of cells contributing to each state. d, Representative genes defining C2 signaling, C1 ECM-remodeling, C0 competence-maintenance and C8 competence-acquisition states. Color denotes relative expression and circle size denotes the percentage of cells in which the gene was detected. e, Inferred communication from C2 signaling fibroblasts to C0, C1 and C8 receiving states across significant ligand–receptor pathways. Circle size represents summed communication probability and color represents −log10(P). f, Leading interpretable C2-derived ligands predicted to act on C0, C1 and C8, grouped by their putative effects on ECM/adhesion, WNT modulation, matrix remodeling and competence-associated programs. These ligand–target relationships are computational predictions rather than evidence of causal signaling. g, Difference in competence-module score between NCAM1-positive and NCAM1-negative cells within the indicated cluster and time point. Positive values indicate higher scores in NCAM1-positive cells; color denotes the score difference. Adjusted significance is shown as *P < 0.05, **P < 0.01 and ***P < 0.001; ns, not significant. h, Summary model in which transient C2 signaling and C1 ECM-remodeling states support C0 competence maintenance and C8 competence acquisition, creating a shared fibroblast niche from which distinct early-DC/dermal-papilla, endothelial-, pericyte- and glial-associated fate-entry routes emerge. Dashed arrows denote candidate transitions inferred from state mapping and not direct lineage relationships. The single-cell RNA-sequencing and communication analyses represent one sequencing dataset (n = 1).

The component states were distinguished by their molecular programs. C2 expressed secreted and matrix-remodeling factors, C1 expressed a contractile and matrix-producing program, C0 combined NCAM1 with competence-maintenance genes, and C8 expressed APCDD1, ROBO2 and other competence-acquisition genes (Fig. 2d and Supplementary Fig. 1b,c). Direct differential-expression comparisons further distinguished C2 as an early and transient signal-provider population enriched for SPP1, FGF7, CXCL12, PDGFA, INHBA, EREG, inflammatory chemokines and matrix metalloproteinases relative to C1/C8 competent fibroblasts (Supplementary Fig. 5a–c). These differences were strongest at PWD12 and PWD13 and diminished by PWD14. The timing, breadth of secreted factors and matrix-remodeling activity identify C2 as a candidate transient signaling organizer that precedes or accompanies expansion of multiple competence- and branch-associated populations, rather than as an obligatory intermediate within any single trajectory.

CellChat analysis predicted extensive communication from C2 toward C0, C1 and C8 through collagen, fibronectin, tenascin, laminin, angiopoietin-like, NCAM, thrombospondin and noncanonical WNT pathway components (Fig. 2e). The leading predicted ligands differed among receiving states: C0 received candidate extracellular-matrix, adhesion and WNT-modulatory inputs; C1 received signals associated with matrix production and contractility; and C8 received matrix and competence-associated inputs (Fig. 2f). Temporal analysis likewise predicted C2-to-C8 communication through NCAM1, SPP1, FGF7, WNT5A and THBS pathways (Supplementary Figs. 1d and 5d). These computations nominate candidate interactions and do not demonstrate that any individual ligand causes competence acquisition.

Within the competence-associated clusters, NCAM1-positive cells generally carried higher competence-module scores than their NCAM1-negative counterparts, particularly in C0 and C1 at the time points shown (Fig. 2g). This within-state comparison argues that NCAM1 marks a transcriptional property of competent cells rather than merely reflecting the increased abundance of an NCAM1-rich cluster. The combined temporal, molecular and communication analyses support a division of labor within the regenerative niche. The combined temporal, molecular and communication analyses support a division of labor within the regenerative niche. C2 is positioned as a candidate transient signaling organizer, C1 is associated with extracellular-matrix remodeling, and C0/C8 maintain or acquire competence before cells populate distinct fate-entry routes (Fig. 2h). In this model, C2 need not serve as the direct precursor of each fate; instead, it may transiently coordinate the states and signaling conditions under which neighboring fibroblasts become competent. The sequencing data support this organizational model but do not establish a causal niche hierarchy.

### Competent fibroblasts enter distinct regenerative fate programs

We next tested whether the competence field was connected to multiple regenerative outcomes. A fibroblast-centered integration of PWD12-PWD14 wounds positioned a broad fibroblast core adjacent to four separable continua: dermal-condensate and dermal-papilla, endothelial, pericyte and glial-associated states (Fig. 3a). Day-resolved projections showed progressive redistribution from fibroblast-core and transition compartments toward branch-associated populations across the three time points (Fig. 3b). Computational stage ordering independently organized each branch from early fibroblast-associated states through transition states toward its corresponding fate program (Fig. 3c). Because these data were sampled cross-sectionally, the trajectories describe transcriptional continua and likely directions of state progression; they do not track conversion of individual cells.

**Figure 3.**
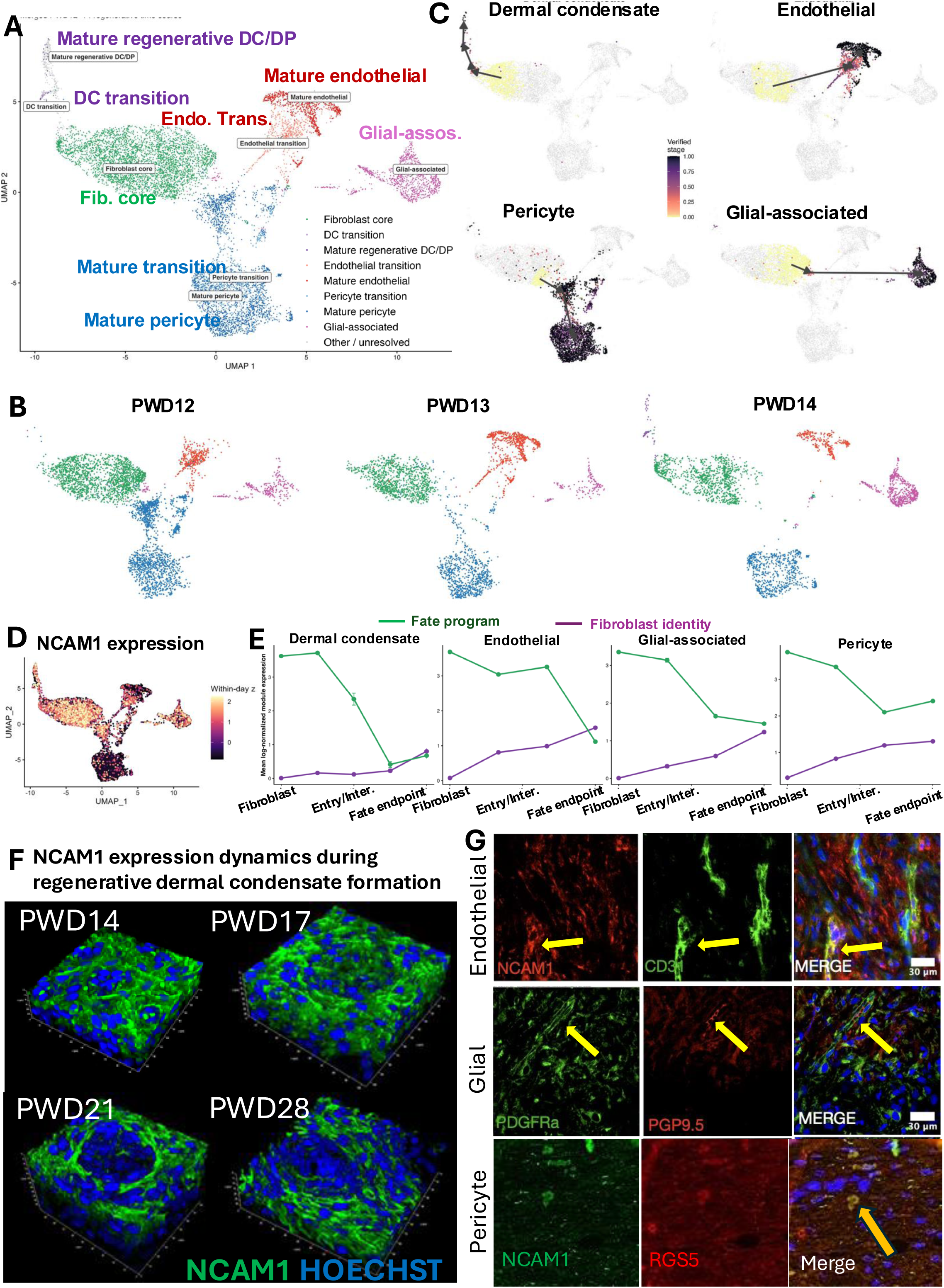
Competent fibroblasts enter distinct dermal-condensate, endothelial, pericyte and glial trajectories. a, Integrated fibroblast-centered UMAP of PWD12–14 wounds showing a fibroblast core and connected DC-transition, mature regenerative DC/dermal-papilla (DP), endothelial-transition, mature endothelial, pericyte-transition, mature pericyte and glial-associated populations. Other or unresolved cells are shown in grey. b, The same UMAP coordinates separated by PWD12, PWD13 and PWD14, illustrating the temporal distribution of fibroblast-core and branch-associated states. c, Inferred stage progression along the DC, endothelial, pericyte and glial-associated trajectories. Color denotes the verified trajectory stage from early (yellow) to late (dark purple), and arrows indicate the direction of computational ordering. These cross-sectional trajectories describe state continua and do not establish direct conversion of individual cells. d, NCAM1 expression projected onto the integrated UMAP. RNA values were normalized and standardized within each day before integration to reduce day-dependent scale effects. e, Mean fibroblast-identity and fate-program module scores across ordered states for each trajectory. Fibroblast identity generally declines as the corresponding DC, endothelial, glial-associated or pericyte program rises; early branch-associated states can retain substantial fibroblast identity. f, Representative three-dimensional immunofluorescence images showing NCAM1 dynamics during regenerative dermal-condensate formation at PWD14, PWD17, PWD21 and PWD28. Nuclei are labeled with Hoechst. g, Representative immunofluorescence images of predicted transitional populations associated with endothelial (NCAM1 and CD31), glial (PDGFRA and UCHL1/PGP9.5) and pericyte (NCAM1 and RGS5) trajectories. Arrows indicate cells or structures with spatially overlapping markers; nuclei are blue. Scale bars, 30 μm where shown. The single-cell RNA-sequencing analysis represents one sequencing dataset (n = 1); staining was performed in three biological samples (n = 3).

Module-score analysis revealed a shared logic across the four branches. Fibroblast identity was highest in source and early-entry states and generally declined as dermal-condensate, endothelial, pericyte or glial-associated programs increased (Fig. 3e and Supplementary Fig. 6a). Importantly, the earliest branch-associated populations retained substantial fibroblast identity. We therefore classified these cells as fate-entry intermediates rather than mature alternative cell types. Heat maps of the underlying genes confirmed staged loss of the shared fibroblast program and acquisition of coherent branch-specific modules, and a crosswalk between source clusters and simplified stages ensured that early intermediates remained within the fibroblast compartment until the alternative program became dominant (Supplementary Fig. 8).

NCAM1 was distributed across the fibroblast core and several transition regions, but its dynamics were not identical across branches (Fig. 3d). Ordered-state analysis showed that NCAM1 expression changed across early-entry, intermediate and mature stages of the four trajectories, consistent with a cross-branch state marker rather than a stable marker of any single terminal fate (Supplementary Fig. 6b). Three-dimensional imaging during dermal-condensate formation similarly showed that NCAM1-positive cells reorganized between PWD14 and PWD28 as condensates and nascent follicular structures emerged (Fig. 3f). Co-staining identified cells with overlapping NCAM1 and CD31 near endothelial-associated structures, PDGFRA and UCHL1/PGP9.5 within glial-associated regions, and NCAM1 and RGS5 along the pericyte-associated route (Fig. 3g). These spatial observations validate the presence of predicted transitional phenotypes without establishing that marker overlap alone represents direct fate conversion. Together, the data support a root-associated competent fibroblast manifold that retains fibroblast identity during early branch entry and subsequently resolves into distinct regenerative programs.

### Shared niche signals accompany branch-specific fate entry

To determine how a common competence field could support different non-appendage outcomes, we compared inferred incoming communication across endothelial-, pericyte- and glial-associated transition populations. At PWD13, C2 signaling, C1 matrix-remodeling and C0/C8 competent fibroblasts formed a densely connected network with all three transition populations (Fig. 4a). Collagen, fibronectin, thrombospondin, laminin and tenascin pathways contributed broadly across receivers, defining a shared physical matrix and adhesion environment (Fig. 4b). Midkine, NCAM, pleiotrophin and noncanonical WNT signals also appeared across more than one branch, whereas other interactions were preferentially associated with individual receivers (Fig. 4b and Supplementary Fig. 7).

**Figure 4.**
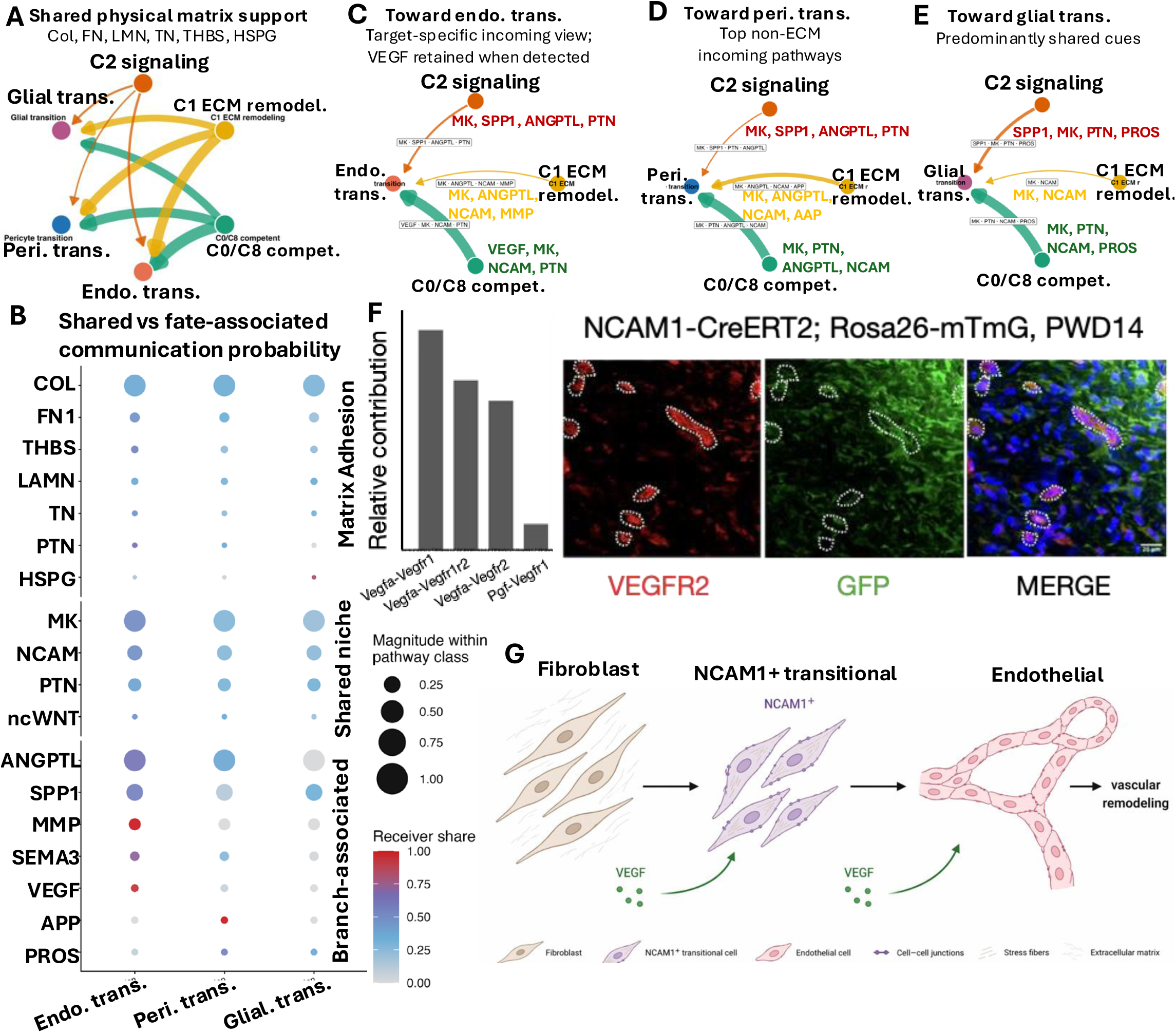
A shared regenerative niche supports distinct endothelial, pericyte and glial-associated fate-entry programs. a, Network representation of inferred matrix and adhesion communication among C2 signaling, C1 ECM-remodeling and C0/C8 competent fibroblasts and endothelial-, pericyte- and glial-associated transition populations from PWD13 scRNA-seq. Edge color denotes the sending population and edge width denotes inferred interaction strength. b, Shared and branch-associated incoming communication programs in endothelial-, pericyte- and glial-associated transition states. Circle size represents pathway magnitude calculated separately within each pathway class, and color denotes the relative receiver share. c–e, Selected signaling routes predicted to converge on endothelial (c), pericyte (d) and glial-associated (e) transition states. Labels indicate representative matrix (MK), SPP1, PTN, ANGPTL, NCAM, MMP, VEGF, APP and PROS interactions retained in the analysis. The diagrams summarize candidate communication routes and do not demonstrate causal induction of fate. f, Relative contributions of VEGF ligand–receptor pairs to the inferred endothelial-associated signal (left) and representative PWD14 wound-center imaging from Ncam1-CreERT2;Rosa26-mTmG mice stained for VEGFR2 (red) and GFP (green) (right). Dashed outlines mark GFP-lineage cells associated with VEGFR2-positive vascular structures. g, Working model in which shared niche-derived signals act on fibroblasts and NCAM1-positive transitional cells during acquisition of an endothelial-associated phenotype and vascular remodeling. The communication analysis represents one single-cell RNA-sequencing dataset (n = 1); staining was performed in three biological samples (n = 3).

The endothelial-associated transition received predicted inputs through matrix, SPP1, ANGPTL, PTN, NCAM and VEGF pathways; the pericyte transition shared several of these inputs but showed a distinct balance that included APP-associated communication; and the glial-associated transition was supported by a partially overlapping set containing matrix, PTN, NCAM and PROS-related interactions (Fig. 4c-e). This organization argues against a single instructive signal that generates all fates. Instead, a common matrix and adhesion niche appears to provide permissive support, upon which branch-associated signals may bias fate entry.

We examined the endothelial route in greater detail because its predicted branch-associated program contained a prominent VEGF component. VEGFA-VEGFR interactions accounted for the largest fractions of the inferred VEGF signal, with additional contributions from other VEGF ligand-receptor pairs (Fig. 4f). In Ncam1-CreERT2;Rosa26-mTmG wounds, GFP-positive lineage cells were incorporated into VEGFR2-positive vascular structures at PWD14 (Fig. 4f). Independent whole-wound communication analysis also identified an NCAM1-positive endothelial-transition population as a strong receiver of VEGF and matrix-associated inputs in large wounds (Supplementary Fig. 9). These findings link NCAM1-positive transitional cells to an angiogenic signaling environment and vascular remodeling (Fig. 4g). They do not establish that VEGF is sufficient to convert fibroblasts into endothelial cells, but they provide a mechanistic framework for testing how a shared niche is resolved into an endothelial-associated program.

### PDGFRA and NCAM1 lineages contribute to regenerative compartments

We further used complementary lineage-tracing strategies to verify the prediction. Pdgfra-CreERT2, Ncam1-CreERT2 and Cdh5-CreERT2 mice were crossed to the Rosa26-mTmG reporter, induced with tamoxifen before wounding and analyzed between PWD14 and PWD28 (Fig. 5a). This design labeled cells expressing each driver before injury and allowed their progeny to be evaluated during regeneration.

**Figure 5.**
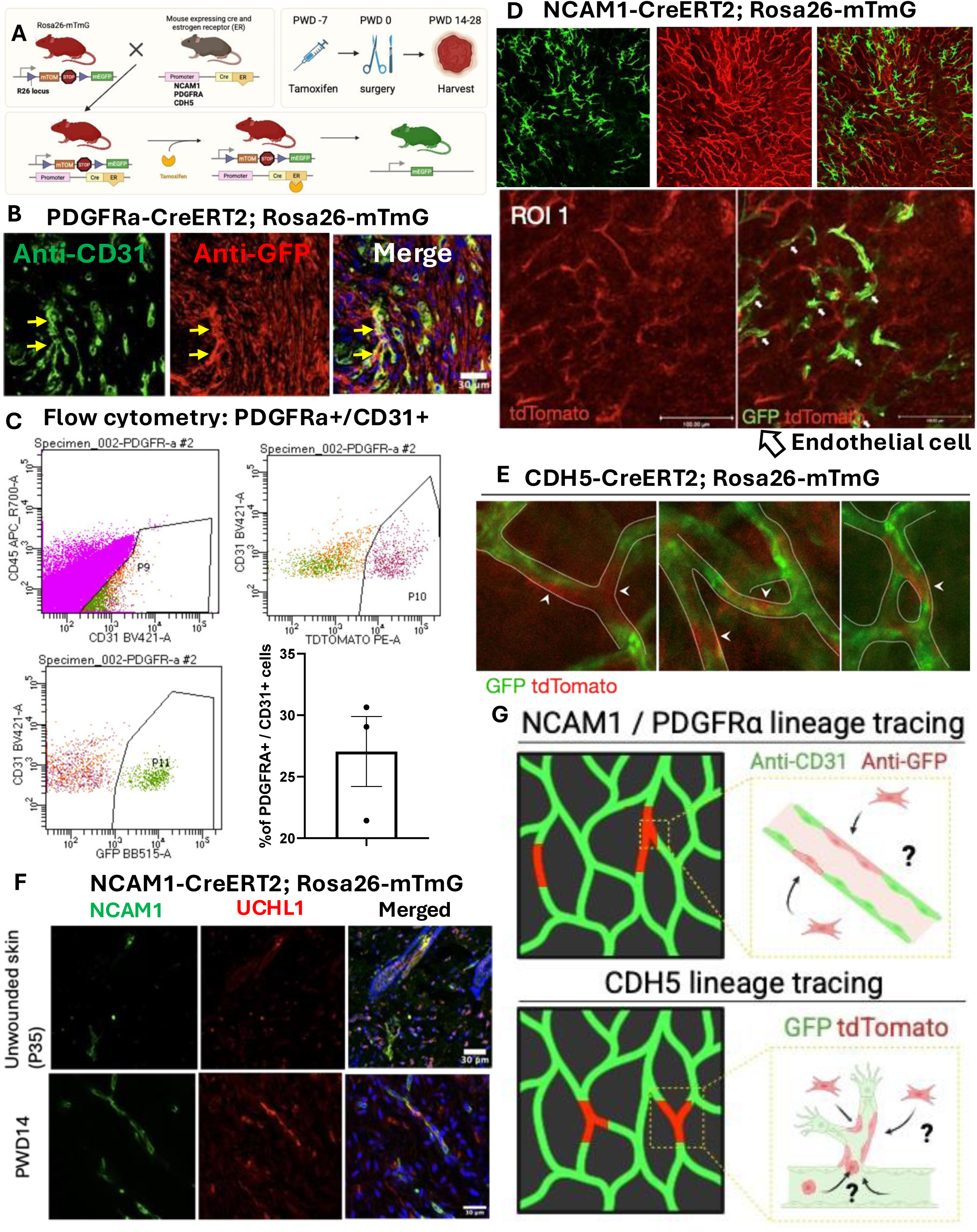
PDGFRA- and NCAM1-lineage cells contribute to multiple regenerative fates. a, Lineage-tracing strategy using Pdgfra-CreERT2, Ncam1-CreERT2 or Cdh5-CreERT2 mice crossed to the Rosa26-mTmG reporter. Tamoxifen was administered 7 days before wounding (PWD−7), wounds were generated at PWD0, and tissues were collected between PWD14 and PWD28. Cre-mediated recombination switches reporter expression from membrane tdTomato to membrane GFP. b, Representative PWD14 wound section from a Pdgfra-CreERT2;Rosa26-mTmG mouse stained for CD31 (green) and GFP (red). Arrows indicate PDGFRA-lineage cells associated with CD31-positive vascular structures. Scale bar, 30 μm. c, Representative flow-cytometry gating used to identify CD31-positive cells carrying the PDGFRA-lineage reporter, together with their quantification. Each point represents one biological sample (n = 3); bars show mean ± SD. Approximately one-quarter to one-third of the analyzed CD31-positive population was reporter-positive under the displayed gating strategy. d, Representative overview and wound-center region of interest from Ncam1-CreERT2;Rosa26-mTmG mice at PWD14. GFP-positive NCAM1-lineage cells are shown against the unrecombined tdTomato-positive tissue background; arrowheads indicate lineage-labeled cells incorporated into or closely associated with vascular structures. e, Representative Cdh5-CreERT2;Rosa26-mTmG images used as a reciprocal endothelial-lineage control. Dashed outlines and arrowheads identify vascular structures and reporter-labeled endothelial cells. f, NCAM1 and UCHL1/PGP9.5 immunofluorescence in uninjured skin (postnatal day 35) and PWD14 wounds from Ncam1-CreERT2;Rosa26-mTmG mice, showing NCAM1-lineage association with nerve/glial structures. Scale bars, 30 μm. g, Schematic comparison of the questions addressed by NCAM1/PDGFRA lineage tracing and reciprocal CDH5 lineage tracing. Staining was performed in three biological samples (n = 3). Together, the data support fibroblast-lineage contribution to vascular- and nerve/glial-associated regenerative populations.

In Pdgfra-CreERT2;Rosa26-mTmG wounds, GFP-positive lineage cells were detected within or closely associated with CD31-positive vascular structures at PWD14 (Fig. 5b). Flow cytometry provided orthogonal support: approximately one-quarter to one-third of the analyzed CD31-positive population carried the PDGFRA-lineage reporter under the displayed gating strategy (Fig. 5c). The presence of reporter-positive cells in this compartment was therefore not limited to isolated histologic events. In Ncam1-CreERT2;Rosa26-mTmG wounds, GFP-positive lineage cells were likewise found throughout the wound center and along vascular structures (Fig. 5d). Cdh5-CreERT2 tracing labeled the expected endothelial vascular network and served as a reciprocal endothelial-lineage comparator (Fig. 5e).

The NCAM1 lineage also extended beyond the vascular compartment. Compared with uninjured skin, PWD14 wounds contained NCAM1-lineage cells associated with UCHL1/PGP9.5-positive nerve or glial structures (Fig. 5f). These data are consistent with the glial-associated trajectory and with the broader distribution of NCAM1 across fate-entry states. Taken together, the lineage experiments demonstrate that pre-wound PDGFRA- and NCAM1-expressing lineages contribute to multiple regenerative compartments, including vascular- and nerve/glial-associated populations (Fig. 5g). In conjunction with the transcriptional continua, the data support fibroblast-lineage contribution during regeneration. They do not establish clonal multipotency or direct conversion of a mature fibroblast into a mature endothelial, pericyte or glial cell.

### NCAM1 perturbation redistributes regenerative tissue outcomes

The preceding analyses identified NCAM1 as a feature of the competence manifold and of several fate-entry routes. We therefore asked whether changing NCAM1 abundance or adhesive function altered regenerative outcome. Cimetidine (Fukuda, Kusama et al. 2008) treatment reduced NCAM1 immunofluorescence in PWD14 wounds in a dose-dependent manner, whereas NCAM1 Fab treatment progressively disrupted the organization of CD31-positive vascular networks at PWD19 (Fig. 6a). Both perturbations also changed the number of regenerated hair follicles assessed at PWD28 (Fig. 6a-c).

**Figure 6.**
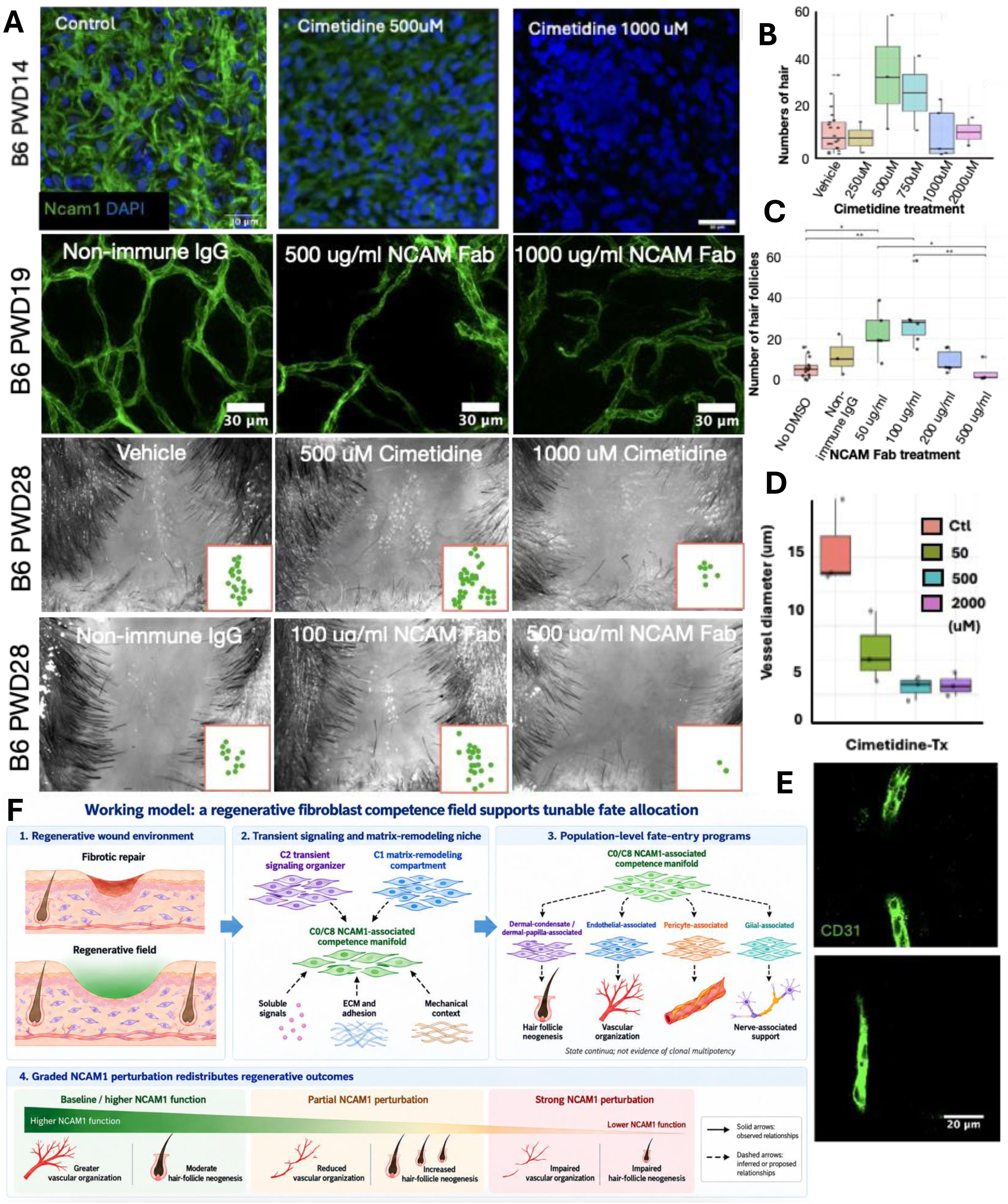
NCAM1 signaling tunes regenerative cell-fate decisions. a, Representative images after pharmacological or antibody-mediated NCAM1 perturbation in C57BL/6 (B6) wounds. Top, NCAM1 immunofluorescence at PWD14 after vehicle, 500 μM cimetidine or 1,000 μM cimetidine treatment; nuclei are stained with DAPI. Scale bar, 10 μm. Second row, CD31-positive vascular networks at PWD19 after nonimmune IgG, 500 μg ml−1 NCAM1 Fab or 1,000 μg ml−1 NCAM1 Fab treatment. Scale bars, 30 μm. Third and fourth rows, representative PWD28 wounds after the indicated cimetidine or NCAM1 Fab treatments; green dots in the insets mark regenerated hair follicles. b, Numbers of regenerated hair follicles following vehicle or graded cimetidine treatment (250–2,000 μM). c, Numbers of regenerated hair follicles following no-DMSO, nonimmune-IgG or graded NCAM1 Fab treatment (50–500 μg ml−1). d, Vessel diameter following control or cimetidine treatment (50, 500 or 2,000 μM). e, Representative CD31-positive vessels in untreated and 750 μM cimetidine-treated wounds. Scale bar, 20 μm. f, Working model of the regenerative fibroblast competence field and quantitative NCAM1-dependent fate allocation. Large regenerative wounds establish a transient niche containing a candidate C2 signaling-organizer population and a complementary C1 matrix-remodeling compartment. These populations are predicted to support a C0/C8 NCAM1-associated competence manifold connected to dermal-condensate/dermal-papilla-, endothelial-, pericyte- and glial-associated fate-entry programs. Dashed arrows denote relationships inferred from temporal state mapping, ligand–receptor analysis or the proposed mechanochemical framework and do not establish clonal multipotency. Graded NCAM1 perturbation produced a non-monotonic redistribution of experimentally measured outcomes: baseline or higher NCAM1 function was associated with greater vascular organization and moderate hair-follicle neogenesis; partial perturbation reduced vascular organization while increasing hair-follicle neogenesis; and stronger perturbation impaired both outcomes. The model supports quantitative allocation among regenerative tissue programs rather than direct conversion between endothelial and appendage-forming cells. Staining was performed in three biological samples (n = 3). Quantified animal studies used six animals per group (n = 6) unless otherwise indicated. For b–d, boxes show the median and interquartile range, whiskers show SD, and statistical comparisons were performed using one-way ANOVA; *P < 0.05 and **P < 0.01.

The response was non-monotonic. Intermediate cimetidine doses increased hair follicle neogenesis relative to vehicle, while the highest dose reduced it (Fig. 6b). NCAM1 Fab produced a similar pattern: partial inhibition increased hair follicle number, whereas stronger inhibition reduced the response (Fig. 6c). In parallel, cimetidine reduced vessel diameter across the tested dose range, and treated wounds contained thinner, less connected CD31-positive structures than untreated wounds (Fig. 6d,e). The concordant results obtained by lowering NCAM1 abundance and interfering with NCAM1 function support NCAM1-dependent regulation rather than a response unique to one reagent.

These dose responses separate two consequences of NCAM1 modulation. High NCAM1 abundance or function was associated with connected vascular networks and moderate follicle neogenesis. Partial inhibition shifted the system into an intermediate range in which vascular connectivity decreased while follicle regeneration increased. Strong inhibition impaired both outcomes (Fig. 6f). Thus, NCAM1 is not simply permissive or inhibitory for regeneration as a whole. Its level or function tunes the distribution of regenerative output among competing or interdependent tissue programs. Because the perturbations were applied to the wound as a population and outcomes were measured at the tissue level, the experiments do not imply direct endothelial-to-hair conversion. Instead, they support the central hypothesis that an NCAM1-associated competence state can be quantitatively modulated to bias fate specification between vascular organization and appendage regeneration.

## Discussion

Adult tissue regeneration requires more than replacement of lost cellular mass: it requires reconstruction of the cellular diversity, spatial organization and functional relationships that existed before injury. Here, we used regenerative and non-regenerative skin wounds to define how such a program may be assembled in adult tissue. Our findings support a model with four connected elements (Fig. 6f). First, large wounds establish a morphogenetically competent fibroblast field containing signaling, matrix-remodeling and competence-associated states that are scarce in small, fibrotically repaired wounds. Second, a transient C2 population emerges as a candidate signaling organizer that coordinates competence-associated states rather than serving as an obligatory precursor of every regenerative lineage. Third, competent fibroblasts progress through branch priming, fate entry and maturation, with shared niche signals establishing competence and branch-associated signals progressively restricting fate. Finally, NCAM1 functions as a quantitative control point: different levels of NCAM1-dependent adhesion and signaling redistribute regenerative output among vascular and appendage-associated programs.

The small-versus-large wound comparison was central to distinguishing regeneration-associated states from cell populations that are common to wound repair. Small wounds re-epithelialize and close, but they do not restore the structures removed by injury. By contrast, large wounds contain a central regenerative field capable of producing new hair follicles and other organized tissue components. Single-cell comparison identified signaling and competence-associated fibroblast states, together with early dermal-condensate, endothelial-, pericyte- and glial-associated populations, that were absent or markedly underrepresented in small wounds. These observations argue that wound size does not merely alter the magnitude of a shared repair response. Instead, large wounds create a qualitatively different cellular ecosystem in which fibroblasts gain access to regenerative states and alternative fate programs. Because the sequencing comparison represents integrated cross-sectional datasets rather than replicated wounds processed in parallel, the relative abundances should be interpreted as a map of candidate regeneration-specific states, not as replicate-level estimates of their frequency. Within this ecosystem, reticular or otherwise inactive fibroblasts provide a useful contrast to the competent fibroblast backbone. The root-associated population retained a strong fibroblast identity while expressing an NCAM1-associated competence program and connecting to several branch-entry states. We therefore interpret this population as competent rather than multipotent: it occupies a transcriptional state from which several regenerative programs are accessible, but the present data do not establish that an individual root cell generates every observed fate. Initial tissue injury and inflammation may help activate fibroblasts and move them away from a quiescent reticular state (Gay, Kwon et al. 2013, Nelson, Reddy et al. 2015, Abbasi, Sinha et al. 2020). Injury activation alone, however, appears insufficient to establish regenerative competence, because small wounds contain abundant injury-responsive cells without forming the same signaling and matrix-remodeling niche. We propose that inflammatory activation creates a permissive starting condition(Gay, Kwon et al. 2013, Nelson, Reddy et al. 2015), whereas additional local signals are required to maintain competence and prime cells for particular regenerative trajectories.

The PWD12-PWD14 analysis further resolves reprogramming into a sequence of states. Competent root-associated fibroblasts retain core fibroblast programs; poised or branch-biased cells begin to express elements of an alternative fate while remaining recognizably fibroblastic; entry and intermediate states progressively lose fibroblast identity; and early and mature states consolidate lineage-specific programs. This staged organization was observed along dermal-condensate/dermal-papilla, endothelial, pericyte and glial-associated trajectories. The results favor a model of progressive restriction: regeneration transiently broadens the range of accessible cell states, after which local signals stabilize one of several lineage-specific regenerative programs. This framework also distinguishes adult regeneration from a literal replay of embryonic development. The wound may reuse developmental regulators, but these factors operate in an inflammatory, mechanically remodeled and heterogeneous adult tissue environment and therefore constitute regeneration-specific programs rather than simple developmental recapitulation.

The regenerative field is supported by a specialized niche rather than by competent cells alone. An important principle emerging from other regenerative systems is that transient intermediate populations can function as signaling organizers. In the hair follicle, transit-amplifying cells do not simply produce differentiated progeny; they become a temporary signaling center that coordinates the behavior of neighboring stem and progenitor compartments (Hsu, Li et al. 2014). The large-wound C2 population may perform an analogous organizational function. C2 appeared transiently before overt tissue patterning and expressed a broad combination of growth factors, inflammatory mediators, matrix-remodeling enzymes and adhesion-associated signals predicted to act on several competence-associated populations. Its inferred role is therefore not necessarily to enter each regenerative trajectory, but to help create the conditions under which neighboring fibroblasts acquire or maintain competence. C1 provided a complementary extracellular-matrix and contractile program. Together, C2 and C1 suggest a division of labor between a transient cellular signaling center and a matrix-remodeling compartment. Communication analysis predicted that these states provide shared inputs to the competent fibroblast backbone and more selective inputs to endothelial-, pericyte- and glial-associated transition states.

The niche should therefore be viewed as a combined biochemical and mechanical signaling system. Soluble ligands can influence receptor activity and transcription, whereas matrix composition, proteolysis and adhesion determine ligand presentation, cell position, force transmission and the duration of cell-cell contact (Bonnans, Chou et al. 2014). In this model, the matrix functions beyond a scaffold surrounding reprogramming cells (Hynes 2009); it is an active component that helps determine which signals can be received and how long a branch-primed state remains reversible. Because CellChat(Jin, Guerrero-Juarez et al. 2021) infers communication from ligand and receptor expression, these networks identify plausible routes of coordination rather than demonstrating that any individual interaction is sufficient to induce fate.

The endothelial-associated branch provides the clearest example of adult fate acquisition in this study. Along the inferred trajectory, fibroblasts retained PDGFRA and extracellular-matrix programs before progressively acquiring KDR/VEGFR2, CDH5, PECAM1 and other endothelial features. NCAM1-positive endothelial-transition cells were predicted to receive VEGF and matrix-associated signals from the surrounding niche. PDGFRA-lineage labeling, NCAM1-lineage imaging and flow cytometry further identified lineage-labeled cells within the CD31-positive compartment, while reciprocal CDH5-lineage analysis provided an endothelial reference. Taken together, these data are consistent with an adult mesenchymal-to-endothelial transition(Ubil, Duan et al. 2014) during regenerative wound healing and extend prior demonstrations that endothelial identity can be induced experimentally in fibroblasts. This is a comparatively uncommon example of fibroblast-lineage contribution to an endothelial-associated population in vivo. Nevertheless, the conclusion should remain bounded by the resolution of the experiments: cross-sectional trajectories do not track individual cells, Cre-based lineage labeling reports ancestry rather than the mechanism of conversion, and cell fusion or labeling of rare non-fibroblast populations cannot be excluded without clonal and orthogonal validation.

NCAM1 is positioned at the intersection of competence, adhesion and fate entry. Homophilic NCAM1-NCAM1 interactions can stabilize contacts between neighboring cells, whereas NCAM1-dependent engagement of FGFR and related signaling pathways can influence migration, cytoskeletal organization and survival(Kiselyov, Soroka et al. 2005). In the wound, NCAM1 was enriched across competence-associated fibroblasts and showed trajectory-dependent changes as cells approached different endpoints. It is therefore unlikely to act as a binary marker or a single lineage determinant. Instead, NCAM1 abundance and engagement may regulate the strength and duration of contact-dependent signaling, the ability of cells to move within the wound bed, and the persistence of a plastic competence state. These functions provide a plausible mechanism by which one adhesion molecule can participate in several stages of regeneration without specifying the same outcome in every cell.

The graded perturbation experiments reveal a tuning relationship rather than a simple requirement for NCAM1. Partial NCAM1 perturbation reduced vessel diameter or vascular connectivity while increasing hair-follicle regeneration, whereas stronger inhibition impaired both vascular organization and appendage regeneration. The magnitude of NCAM1-dependent adhesion or signaling therefore appears to determine which outcomes remain accessible to the competent population. One interpretation is that sustained NCAM1 activity favors endothelial entry and vascular organization, whereas an intermediate level preserves competence but reduces vascular commitment, increasing the relative opportunity for dermal-condensate specification. Stronger inhibition may cross a second threshold at which cell cohesion, migration or competence maintenance also fails. NCAM1 consequently behaves less like an on–off fate determinant than a quantitative control point that redistributes population-level regenerative output. This interpretation explains the increased hair regeneration following partial inhibition without requiring direct endothelial-to-dermal-condensate conversion.

Quantitative control may be a general principle of regenerative systems. Hair-follicle populations respond differently according to injury density, selective modulation of gp130 pathway output can enhance regeneration, and wound-induced hair follicle neogenesis occurs within an intermediate mechanical range rather than varying monotonically with tissue stiffness(Chen, Wang et al. 2015, Harn, Wang et al. 2021, Harn, Chiu et al. 2022). The NCAM1 dose response extends this principle to cell adhesion and fate determination. Our previous work showed that pharmacological softening expands the morphogenetically competent wound region, whereas the present findings suggest a cellular interface through which mechanical context could influence the resolution of competence. Consistent with the ability of matrix elasticity to direct lineage specification(Engler, Sen et al. 2006), changes in stiffness or contractility could alter NCAM1 engagement, cell-cell contact, growth-factor responsiveness and nuclear mechanotransduction, thereby changing the stability of competent and branch-committed states. This model does not establish NCAM1 as a direct mechanosensor; it positions NCAM1 within a mechanochemical system that converts quantitative changes in the wound environment into different probabilities of fate entry.

Several questions follow from this model. First, the upstream signals that move inactive dermal fibroblasts into the competent state remain unresolved. Inflammatory cues are likely contributors, but comparison with small wounds indicates that additional spatially restricted matrix, adhesion and growth-factor inputs are required. Second, the molecular basis by which quantitative niche inputs are converted into fate restriction should be tested at the chromatin level. We predict that root-associated competent cells transiently maintain accessibility across distal cis-regulatory elements linked to several regenerative programs. As cells become poised and enter a branch, graded changes in mechanical context, NCAM1 engagement and receptor signaling may redistribute accessibility across these regulatory domains, stabilizing one program while closing alternatives. Matched scRNA-seq and scATAC-seq across defined NCAM1 perturbation and matrix-stiffness conditions will be required to determine whether distal CRE accessibility changes with input magnitude and contributes causally to branch choice. Such experiments should also reveal whether partial NCAM1 perturbation favors dermal-condensate specification by altering competence maintenance, selectively weakening endothelial entry, or both.

The study has additional limitations. The integrated single-cell analysis represents one sequencing dataset assembled from different studies and post-wounding days, so day, platform and biological condition cannot be fully separated. Pseudotime provides complementary evidence for state ordering but do not prove lineage direction. CellChat identifies candidate communication routes rather than causal signals. PDGFRA- and NCAM1-lineage tracing supports fibroblast-lineage contribution, but higher-resolution clonal tracing and reciprocal perturbation will be required to establish conversion efficiency and exclude alternative explanations. The pericyte and glial-associated branches are supported by transcriptional continua and spatial marker overlap but remain less functionally validated than the endothelial branch. Finally, the relationship between mechanics, NCAM1 abundance and fate specification is supported by convergent observations across experiments; direct demonstration of a causal mechanical-NCAM1 pathway will require controlled manipulation of wound stiffness with matched measurements of lineage outcome.

Together, these findings shift the conceptual boundary between fibrotic repair and regeneration. A regenerative wound does not simply contain more proliferating cells or stronger growth-factor activity. It establishes a transient morphogenetic competence field in which a signal-rich organizer population and a matrix-remodeling compartment support NCAM1-associated competent fibroblasts. These cells subsequently become poised for alternative fates, while branch-associated interactions progressively stabilize endothelial, mural, glial-associated or appendage-forming programs. NCAM1 tuning changes the balance among these outcomes, demonstrating that regenerative fate specifications depends on the magnitude of signaling and adhesion rather than on pathway activation alone. We propose that biochemical signaling, matrix mechanics and cell adhesion converge on regulatory programs that establish, maintain and resolve competence. Defining these quantitative control points may provide a route for shifting adult wounds away from scar-dominated repair and toward organized, multi-structure regeneration.

## Materials and Methods

### Study design

The experimental strategy combined single-cell transcriptomic integration, computational state and trajectory analysis, immunofluorescence, inducible lineage tracing, flow cytometry, and graded pharmacological or antibody-mediated perturbation of NCAM1. Single-cell RNA-sequencing comparisons represent one integrated sequencing dataset (n = 1) and were therefore interpreted descriptively at the dataset level. Immunostaining experiments used three biological samples (n = 3), and quantified animal studies used six animals per group (n = 6) unless otherwise indicated.

### Animals and ethical approval

All animal procedures were performed under a protocol approved by the Institutional Animal Care and Use Committee of the University of Southern California and in accordance with institutional guidelines. C57BL/6J mice were used for wound and perturbation experiments. Mice were housed in climate-controlled facilities under a 12-h light/12-h dark cycle, at 21-26 °C and 30-70% relative humidity, with food and water available ad libitum. Both sexes were included where available.

### Mouse wound models

5-6-week-old C57Bl6 mice of both sexes were used. Dorsal hair was removed and full-thickness excisional wounds extending through the panniculus carnosus were generated under anesthesia. Small repair wounds measured 0.6 × 0.6 cm, whereas large regenerative wounds measured 1.5 × 1.5 cm. Wounds were allowed to heal by secondary intention and were collected at the indicated post-wounding day (PWD). Wound tissues were harvested from the center or from the entire wound bed as specified for each experiment. The general surgical and post-operative procedures followed previously published laboratory protocols (Harn, Wang et al. 2021), with wound dimensions modified as shown in Fig. 1a.

### Lineage tracing

For lineage tracing, Ncam1-P2A-CreERT2 mice (GemPharmatech; heterozygous; C57BL/6J background), Pdgfra-CreERT2 mice (The Jackson Laboratory, stock no. 032770; heterozygous; C57BL/6J background), and Cdh5(PAC)-CreERT2 mice (Tg(Cdh5-cre/ERT2)1Rha; originally generated in the laboratory of Ralf H. Adams and provided by Dr. Chen Yuan Kam, University of California, Los Angeles) were crossed to Rosa26-mTmG reporter mice (The Jackson Laboratory, stock no. 007676). The Cdh5(PAC)-CreERT2 line was used as an inducible endothelial-lineage comparator, following the lineage-labeling framework described by Kam et al. (Kam, Singh et al. 2023).

### NCAM1 perturbation

NCAM1 abundance or function was perturbed during the regenerative patterning window by topical application of cimetidine or an NCAM1 Fab fragment. Reagents were applied directly to the wound surface from PWD10 through PWD14; vehicle-treated wounds served as controls. Cimetidine was tested at 50-2,000 μM across the imaging and dose-response experiments. NCAM1 Fab was tested at 50-1,000 μg ml−1; nonimmune IgG served as the antibody control. The vehicle contained 1% DMSO where applicable. Wounds were collected at PWD14 for NCAM1 staining, at PWD19 for vascular analysis, and at PWD28 for regenerated-hair-follicle counts.

### Genotyping

Genomic DNA was isolated from ear or tail biopsies and amplified by PCR. For the Ncam1-P2A-CreERT2 allele, primers T057354-F1/R1 amplified a 420-bp targeted product and primers T057354-F2/R2 amplified a 321-bp wild-type product. Rosa26-mTmG and Pdgfra-CreERT2 alleles were genotyped using the supplier-recommended protocols. PCR products were resolved by agarose-gel electrophoresis and genotypes assigned from the expected band patterns.

### Tamoxifen induction and lineage tracing

CreERT2-mediated recombination was induced by intraperitoneal tamoxifen administration at 75 mg kg−1 once daily for five consecutive days. The induction course was completed at least 7 days before wounding. Full-thickness wounds were then generated and collected between PWD14 and PWD28. Cre-mediated recombination switched the Rosa26-mTmG reporter from membrane tdTomato to membrane GFP. Endogenous reporter fluorescence was visualized directly and, where indicated, amplified with anti-GFP antibody. Sections and whole mounts were co-stained with CD31, VEGFR2, UCHL1/PGP9.5 or other lineage-associated markers to determine whether reporter-positive cells were incorporated into or closely associated with regenerative structures. These experiments were interpreted as lineage contribution and not as clonal proof of direct transdifferentiation or multipotency. Tamoxifen is dissolved in corn oil at 20 mg/ml.

### Tissue collection and cryosection preparation

Wounded and unwounded dorsal skin was harvested at the indicated time points, rinsed in cold phosphate-buffered saline (PBS), and fixed in 4% paraformaldehyde overnight. For cryosectioning, samples were cryoprotected in sucrose, embedded in optimal cutting temperature compound and sectioned on a cryostat. Sections were mounted on positively charged glass slides and air-dried before staining.

### Whole-mount preparation

For whole-mount analysis, wound tissues were trimmed to remove excess subcutaneous tissue, flattened to expose the wound surface and fixed in 4% paraformaldehyde in PBS for 4-6 h at room temperature. Tissues were washed in PBS and blocked overnight at 37 °C in PBS containing 0.2% Triton X-100, 5% normal donkey serum and 1% bovine serum albumin. Samples were incubated with primary antibodies for 48-72 h and secondary antibodies for 24 h at 37 °C on a rocker, washed and mounted in Vectashield antifade mounting medium (Vector Laboratories). This procedure followed the regenerative-wound whole-mount protocol described previously (Harn, Wang et al. 2021).

### Immunofluorescence staining

Cryosections were rehydrated in PBS, permeabilized in detergent-containing buffer and blocked in serum-containing buffer. Sections were incubated with primary antibodies overnight at 4 °C, washed, and incubated with species-appropriate Alexa Fluor-conjugated secondary antibodies. Whole mounts were permeabilized under conditions optimized for thick tissue, blocked, incubated with primary antibodies for an extended period at 4 °C with gentle rocking, washed extensively and incubated with secondary antibodies. Unless otherwise indicated, primary antibodies were used at 1:50 and secondary antibodies at 1:1000. Nuclei were counterstained with Hoechst 33342 at 1:1,000 or with DAPI, and samples were mounted in antifade medium. The staining workflow was adapted from the previously published section and whole-mount protocols (Harn, Wang et al. 2021, Harn, Jiang et al. 2026).

### Primary antibodies and detection reagents

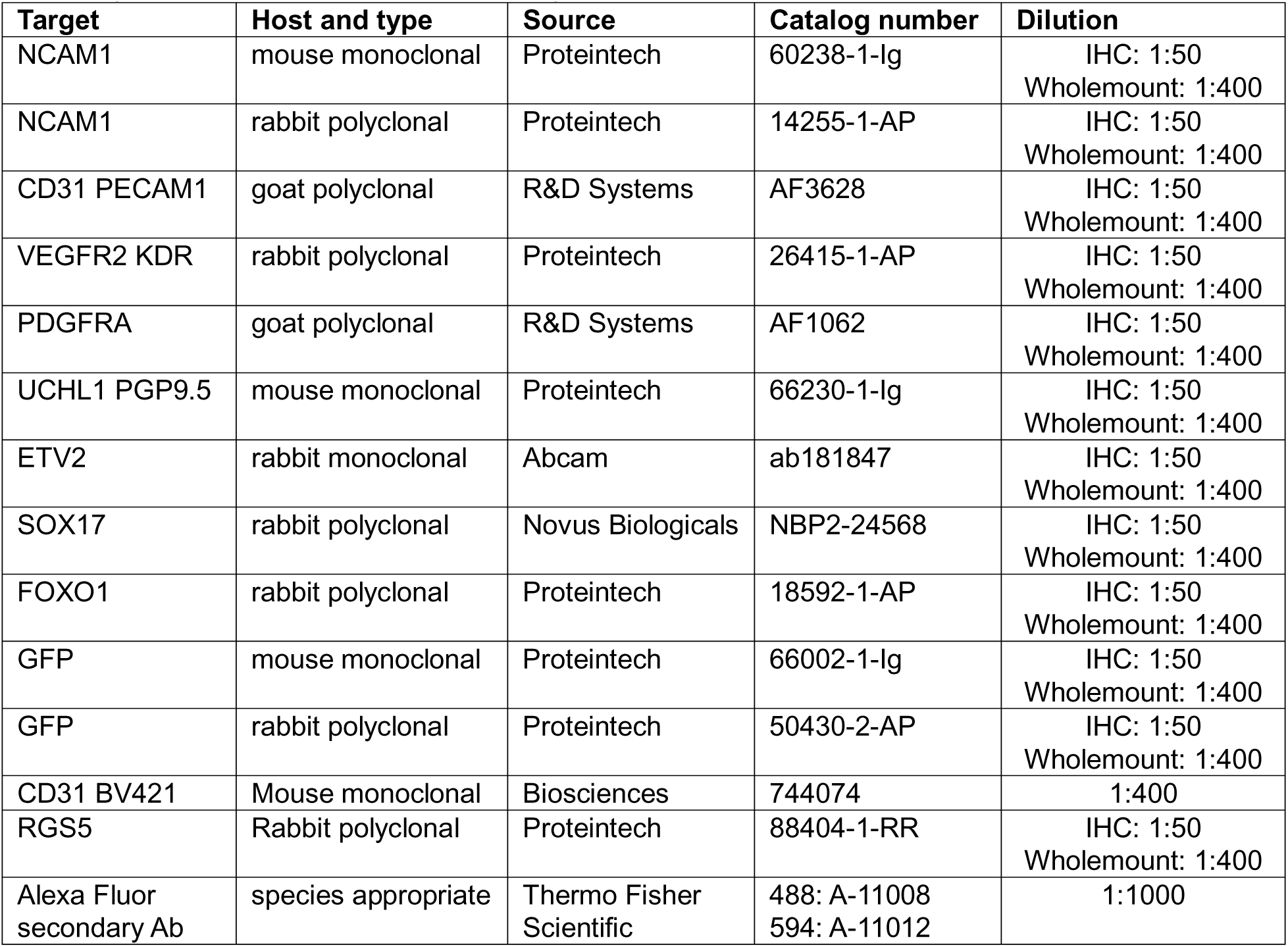

#### Fluorescence imaging and three-dimensional reconstruction

Whole-mount immunostaining imaging was done using a confocal microscope (Stellaris 5, Leica) using 10× air or 63x oil lens. Z-stacks of at least 50 um were taken to reconstruct the 3D structure. Vascular and lineage-labeling image stacks were acquired following the skin-imaging workflow of Kam et al. (Kam, Singh et al. 2023) using a LaVision TriM Scope II laser-scanning microscope equipped with Chameleon Vision II and Chameleon Discovery two-photon lasers. Excitation wavelengths of 940 nm and 1,120 nm were used for live and whole-mount fluorescence imaging. Z-stacks spanning 100-200 μm were acquired at 1-2 μm intervals through Nikon 25×/1.0 or 40×/1.15 water-immersion objectives. Fields of view were 0.44 × 0.44 mm or 0.28 × 0.28 mm, respectively, and larger regions were assembled from sequential tiled fields. Raw stacks were imported into Fiji/ImageJ v1.52p or Imaris v9.5.1; individual optical planes and summed or maximum-intensity projections were used for visualization, and tiled fields were assembled with the Fiji Grid/Collection Stitching plugin(Preibisch, Saalfeld et al. 2009).

### Image analysis

Images were analyzed in Fiji/ImageJ(Schindelin, Arganda-Carreras et al. 2012). Measurements included NCAM1-positive area and fluorescence intensity, marker overlap within defined regions of interest, vessel diameter and vascular organization, and the number of regenerated hair follicles. The same analysis settings were applied to images within each experiment. Three-dimensional images were used to assess the spatial relationship of lineage-labeled cells to vascular, nerve/glial and dermal-condensate structures.

### Regenerated hair follicle quantification

Regenerated hair follicles were quantified from PWD28 large wounds. Follicles within the wound bed were identified from whole-wound images and marked during counting; the total number per wound was used as the biological replicate.

### Flow cytometry of lineage-labeled vascular cells

Pdgfra-CreERT2;Rosa26-mTmG mice were induced before wounding as described above, and wounds were harvested on PWD14. Wound tissue was processed to a single-cell suspension using a workflow adapted from Chen et al. (Chen, Jea et al. 2026). Briefly, excised tissue was trimmed to remove subcutaneous fat and muscle, placed dermis-side down in 0.25% trypsin-EDTA (Sigma, T4049), and incubated overnight at 4 °C. The epidermis was removed with forceps, and the remaining wound tissue was mechanically dissociated in cold PBS. The suspension was neutralized with medium containing 0.05 mM Ca2+, passed sequentially through 70-μm and 40-μm cell strainers (Falcon, 352350 and 352340), and centrifuged at 300 × g for 5 min at 4 °C. Cell pellets were resuspended in staining buffer consisting of 2% Chelex-treated serum without calcium in PBS.

Cells were stained with a BV421-conjugated antibody against CD31/PECAM1 to identify endothelial cells. PDGFRA-lineage cells were identified by Cre-dependent membrane GFP fluorescence from the Rosa26-mTmG reporter; no antibody was required for primary lineage assignment. Nonviable cells were excluded with DRAQ7, following the reference protocol.

Events were gated sequentially to remove debris by forward- and side-scatter, exclude doublets by pulse geometry, and retain viable single cells. CD31-positive events were then defined using unstained, fluorescence-minus-one and/or single-color controls and subdivided by GFP reporter status. The primary outcome was the percentage of CD31-positive endothelial cells derived from the PDGFRA lineage, calculated as 100 × (CD31-positive GFP-positive viable singlets)/(all CD31-positive viable singlets). Data were acquired using cytometer (BD FACS Aria IIu) and analyzed in FlowJo version (v11.2). Single-color compensation controls and appropriate reporter-negative littermate controls were used to establish compensation and positivity gates.

### Single-cell RNA sequencing datasets

The integrated analysis combined public and newly generated mouse wound scRNA-seq datasets. PWD12 large-wound cells were obtained from GEO accession GSE113854 (sample GSM3121363 (Guerrero-Juarez, Dedhia et al. 2019)). The published reads were aligned to mm10 with Cell Ranger v2.1.0. PWD14 data were obtained from the Hic1-lineage wound datasets in GSE108677 (Abbasi, Sinha et al. 2020), distributed within SuperSeries GSE155678; the analyzed libraries included large-wound center and periphery, tdTomato-positive and tdTomato-negative fractions, as applicable. We generated and published the PWD13 data previously (Shkhyan, Flynn et al. 2023), accessible in GEO accession GSM5726293.

The small-wound comparator was obtained from the integrated scarring-wound analysis reported by Phan et al.(Phan, Sinha et al. 2021). Its three constituent small-wound source datasets were merged and treated as one small-wound reference rather than as biological replicates. The Phan et al. dataset (GSE153596)(Phan, Sinha et al. 2021) was generated from 2-mm wounds collected 7 days after injury from P21 mice. The percentages of cells or states were treated as descriptive composition estimates rather than biological-replicate statistics.

### Wound dissociation and single-cell library preparation

For the newly generated scRNA dataset, single cells were isolated from excised large-wound tissue by collagenase/dispase digestion. Cells were stained with antibodies against CD45, CD31 and Ter119 to exclude hematopoietic, endothelial and erythroid cells, respectively, and DAPI was used to exclude nonviable cells. At least 100,000 viable CD45-negative CD31-negative Ter119-negative cells were collected for single-cell capture.

Single-cell capture and droplet generation followed Drop-seq(Macosko, Basu et al. 2015) protocol v3.1 and the original Drop-seq method. Cells at 150,000 cells ml−1, barcoded beads at 175,000 beads ml−1 and droplet-generation oil were co-flowed through a PDMS microfluidic device at 4, 4 and 15 ml h−1, respectively. After droplet breakage, reverse transcription was performed, cDNA was PCR-amplified and magnetically purified, and sequencing libraries were generated by tagmentation and cleaned before sequencing on an Illumina HiSeq 4000 or NovaSeq instrument.

### Raw-read processing and data deposition

The initial scRNA sequencing output was processed in Partek Flow to generate the gene-by-cell expression matrix. Cells with mitochondrial reads comprising 5% or more of total reads were excluded. The resulting expression object was subsequently reanalyzed using the more recent Seurat(Hao, Stuart et al. 2024), module-scoring, trajectory and cell-cell communication workflows described below.

### Single-cell quality control, normalization and integration

Count matrices were analyzed in R using Seurat v5.3.0(Hao, Stuart et al. 2024). Cells with low detected-gene counts, high mitochondrial-transcript fractions or evidence of doublets were removed. Counts were log-normalized, highly variable genes were identified, and scaled expression values were subjected to principal-component analysis. Datasets were integrated using Seurat anchor-based integration to reduce dataset-associated effects. A shared-nearest-neighbor graph was constructed from the integrated principal components; unsupervised clusters were visualized by Uniform Manifold Approximation and Projection.

Whole-wound clusters were annotated conservatively from canonical lineage markers and retained as fibroblast, endothelial, pericyte, immune, epithelial, proliferating or unresolved/mixed populations. The fibroblast compartment was subsetted and reclustered at resolution 0.9. Integrated fibroblast clusters were assigned to inactive/root-like, competence-associated, signaling/remodeling or other specialized states using marker expression, source-cluster crosswalks and post hoc module scores. Percentages were calculated within the retained cells of the relevant condition or within the fibroblast compartment, as stated in the figure legends.

### Differential expression and cluster annotation

Differential expression was tested at the cell level using the Wilcoxon rank-sum test implemented in Seurat, with multiple-testing adjustment by the Benjamini-Hochberg method. Comparisons included C2 signaling fibroblasts versus C1/C8 competent fibroblasts, the large-wound remodeling population versus all other retained fibroblast clusters, and competence-associated C0/C1/C8 states versus inactive C6/C10 states. Genes were ranked by average log2 fold change, adjusted P value and detection fraction. Cell-level adjusted P values were used for marker discovery and were not interpreted as biological-replicate inference.

### Gene modules and NCAM1 stratification

The inactive/root module contained Wnt2, Pi16, Inmt, Cxcl12, Dpep1 and Col14a1. The competence-acquisition module contained Ncam1, Pim1, Ptn, Mdk, Crabp1, Mmp11, Apcdd1, Robo2, Plxna4 and Stra6. The signaling/remodeling module contained Spp1, Il11, Inhba, Mmp3, Mmp9, Mmp10, Saa3 and Hmga2. For branch analysis, the shared fibroblast-identity module contained Pdgfra, Col1a1, Col1a2, Col3a1, Dcn, Lum, Col6a1 and Col6a2. Fate-acquisition modules were: dermal condensate/dermal papilla (Lef1, Sox2, Fgf10, Bmp4, Rspo1, Wif1, Corin and Alpl); endothelial (Kdr, Klf2, Esam, Emcn, Cdh5, Pecam1, Erg and Egfl7); pericyte (Rgs5, Cspg4, Mcam, Pdgfrb, Notch3, Des, Acta2 and Cnn1); and glial-associated (Ngfr, Fabp7, S100b, Sox10, Plp1, Mpz, Mbp and Uchl1). NCAM1 was analyzed separately as a single-gene, cross-branch readout and was not averaged into the fate modules.

Module scores were recalculated directly from normalized RNA expression using case-insensitive matching to the same gene sets in every dataset. A coverage audit confirmed finite scores for all programs and days. Both raw per-cell module means and within-day standardized scores were retained; the latter were calculated separately within each day before integration to reduce day-dependent scale effects in shared visualizations. Within-cluster comparisons contrasted competence-module scores between NCAM1-positive and NCAM1-negative cells.

### Trajectory inference and pseudotime analysis

The final four-fate analysis used the verified analysis backbone stored in verified_multifate_analysis_backbone.rds, including the analysis_identity annotations and the verified_multifate_umap/locked_parent_umap coordinates. Four trajectories were fitted independently with Slingshot(Street, Risso et al. 2018) rather than forcing one universal root. Only regenerative PWD12-PWD14 cells were used; mature or pre-existing small-wound dermal papillae, lymphatic, cycling and contaminating populations were excluded. Early branch-associated cells were retained within the fibroblast compartment until they showed coherent loss of the shared fibroblast module and acquisition of the alternative fate module. The dermal-condensate trajectory followed G8 fibroblasts through A9_s5 branching fibroblasts to early, intermediate and mature regenerative A9_s3 states. The endothelial trajectory followed C1-proximal fibroblasts through G15 entry and C20 intermediate states to mature C9/A15 endothelial cells. The pericyte trajectory followed branch-proximal competent fibroblasts through C23 entry and G2 intermediate states to mature G9/C5/A8 pericytes. The glial-associated trajectory followed G0 glial-biased fibroblasts through entry and intermediate states to repair-associated and myelinating glial endpoints. G8, A9_s5, C1-proximal and G0 cells remained classified as fibroblasts; branch colors began only at verified fate-entry states. The resulting order describes transcriptional continua and not tracked conversion of individual cells.

An earlier PWD13-focused sensitivity analysis used Monocle 3(Cao, Spielmann et al. 2019) on the Seurat v5 object WIHNCTRLchrgtftop.filtered.umap.res.obj.Miller_copy.RDS. RNA counts from resolution-0.9 clusters C1, C3, C5, C9, C13, C16, C18 and C20 were log-normalized, reduced by 50-component PCA and UMAP, and converted to a cell_data_set. Principal graphs were learned with learn_graph using partitioned and unpartitioned fits and a minimal branch length of 10. The dominant graph node nearest C1 was used as the root for order_cells. This analysis was used to assess candidate topology; the final displayed trajectories were the independently fitted, branch-specific Slingshot(Street, Risso et al. 2018) models described above.

### Cell-cell communication analysis

Ligand-receptor communication was inferred with CellChat (Jin, Guerrero-Juarez et al. 2021) using mouse ligand-receptor annotations. The workflow used subsetData followed by identifyOverExpressedGenes with do.fast = TRUE and the presto dependency, then identified overexpressed interactions, calculated communication probabilities, summarized pathway-level probabilities and aggregated the network. Analyses were performed on the relevant integrated fibroblast states and fate-entry populations, with the PWD13 dataset used for the shared-niche and branch-associated networks in Fig. 4. C2 signaling fibroblasts were specified as senders and C0, C1, C8 or endothelial-, pericyte- and glial-associated transition states as receivers, as appropriate. Communication probabilities were summarized by ligand-receptor pair and pathway; pathway-level values were obtained by summing the relevant inferred probabilities. Receiver share was calculated within each pathway class for visualization. Candidate interactions were filtered for statistical support and biological interpretability, with emphasis on matrix/adhesion, SPP1, PTN, ANGPTL, NCAM, MMP, VEGF, APP and PROS signaling. CellChat results were treated as hypothesis-generating predictions and not as evidence of causal fate induction.

### Statistical analysis

Biological replicate numbers are stated in the figure legends. Unless otherwise indicated, staining experiments used n = 3 independent animals and quantified animal studies used n = 6 animals per group. Multi-group comparisons were analyzed by the Kruskal-Wallis test followed by Dunn’s post hoc test with Benjamini-Hochberg correction. All tests were two-sided. Adjusted P < 0.05 was considered significant. Single-cell differential-expression tests used cells as observational units for marker discovery and were not presented as replicate-level hypothesis tests. Box plots show the median and interquartile range; Analyses and plots were generated in R version (4.6.1).

## Acknowledgements and funding

This work is supported by NIH R37 AR 060306.

## Author contributions

CHH: Conceptualization; Data curation; Formal analysis; Validation; writing. CKC: Software, formal analysis. CYK, BVH, AS: Data curation, formal analysis. YCH: Conceptualisation, review and editing. DE: Resources, funding acquisition. CMC: conceptualisation, resources, supervision, funding acquisition, investigation, review and editing. HIH: conceptualisation, data curation, formal analysis, validation, investigation, writing

## Data and code availability

Published PWD12 scRNA-seq data are available from GEO under GSE113854. Published PWD13 scRNA-seq data are available from GEO under GSM5726293. Published PWD14 Hic1-lineage data are available under GSE108677 and SuperSeries GSE155678. The three-source small-wound framework described by Phan et al. is derived from GSE108677, GSE112671 and GSE153596.

