## Supplementary Figures and Tables for "A regenerative fibroblast competence field supports multiple cell fates in adult skin regeneration"

### SI Fig 1. Integrated fibroblast-state analysis identifies competence-associated and signaling populations enriched in large wounds

A

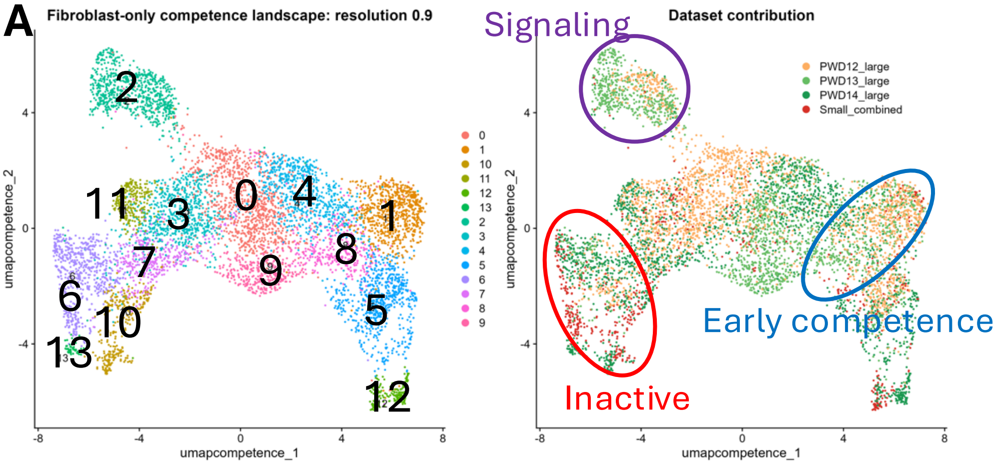


B

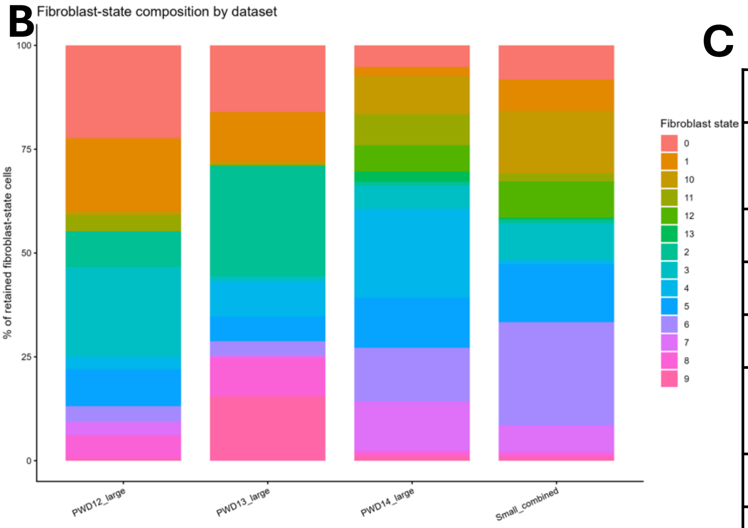


| Cl | Proposed identity | Small | Large |
|---|---|---|---|
| 0 | NCAM1 activated fibroblast | 8.23% | 16.04% |
| 1 | HTRA1/CRABP1 early competent fibroblast | 7.49% | 12.00% |
| 2 | Competence signaling hub | 0.90% | 26.62% |
| 3 | PTN/MDK trophic fibroblast | 8.83% | 1.15% |
| 4 | ASPN/CDH3 matrix-organizing fibroblast | 1.05% | 8.54% |
| 5 | IGFBP2/CYP26B1 G8/S2-related fibroblast | 13.92% | 5.92% |
| 6 | PI16/CLEC3B inactive root-like fibroblast | 24.85% | 3.41% |
| 7 | DLK1/NOV adventitial ECM fibroblast | 6.59% | 0.52% |
| 8 | TWIST/POSTN advanced competent fibroblast | 1.05% | 9.33% |
| 9 | POSTN/CCN4/INHBA activated fibroblast | 0.90% | 15.51% |
| 10 | CLEC3B/INMT reticular quiescent fibroblast | 15.12% | 0.63% |
| 11 | SCX/TNMD tendon-like ECM fibroblast | 1.95% | 0.21% |
| 12 | ALDH3A1/CCL19 specialized fibroblast | 8.68% | 0.10% |
| 13 | SMPD3/PGR rare specialized fibroblast | 0.45% | 0.00% |

C Identifying competence-acquisition program via PWD12 C1/8 vs 6/10 comparison

| Gene | Potential function |
|---|---|
| Crabp1 | Papillary, retinoid-response/state plasticity |
| Htra1 | ECM and growth-factor regulation |
| Mmp11 | Matrix remodeling |
| Igfbp2 | Growth-factor availability and signaling |
| Ptk7 | Noncanonical WNT/planar-polarity signaling |
| Tcf4 | Transcriptional-state regulation |
| Megf6 | Matrix-associated signaling |
| Cpe | Processing of signaling peptides |

D Unique cluster 2: unique signaling hub to C8

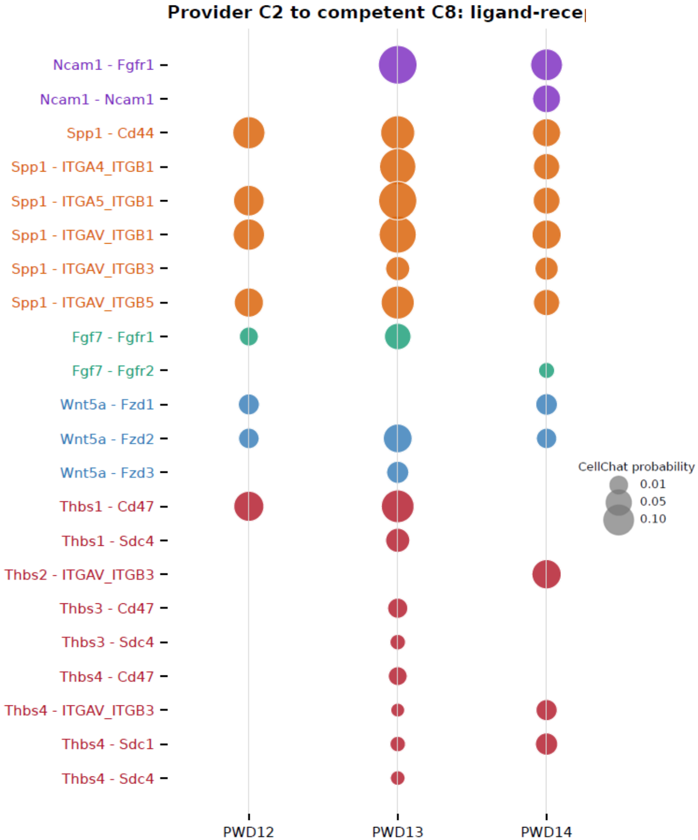

**SI Fig 2. Marker analysis distinguishes fibroblast states between small and large wounds**

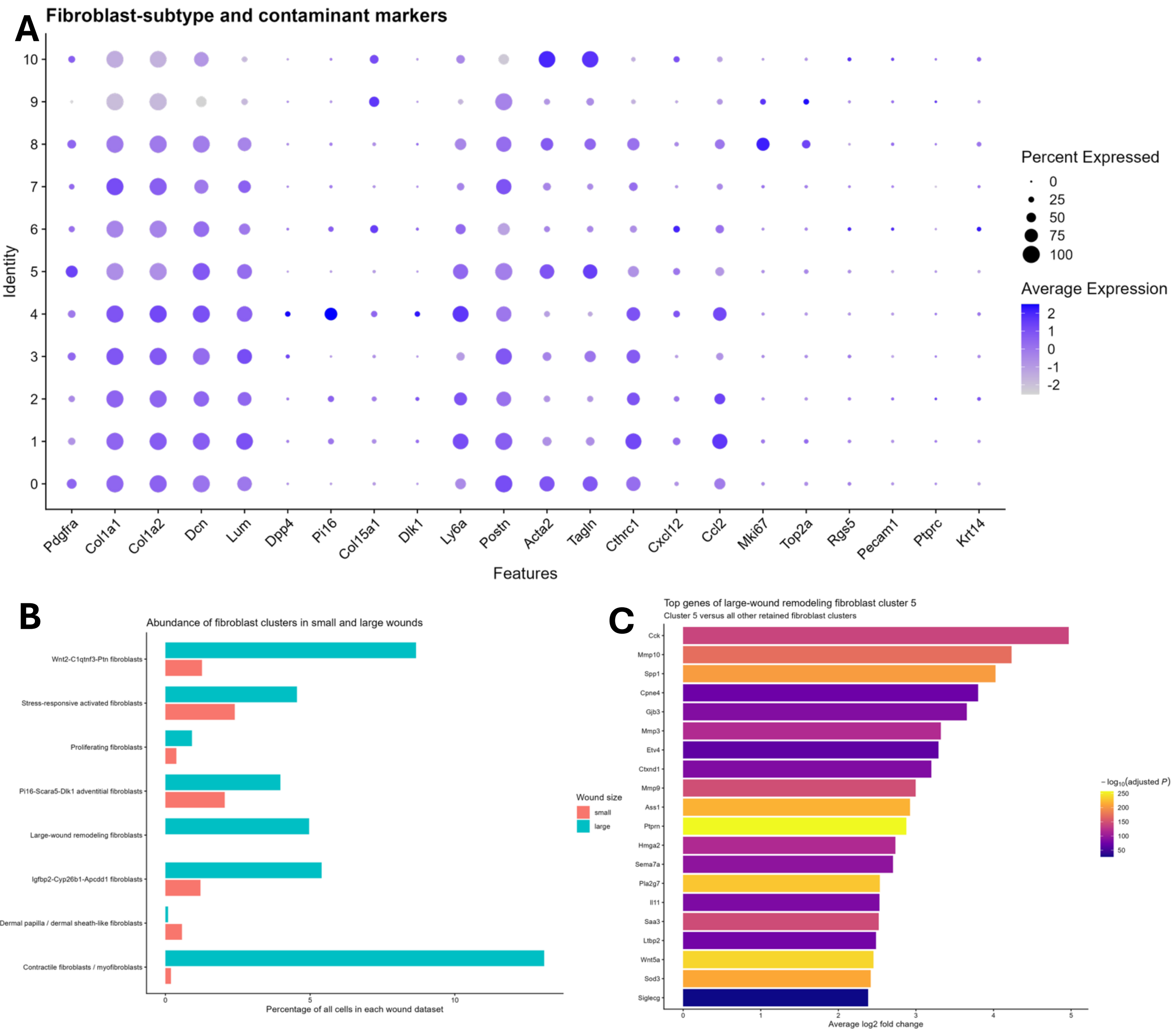


### D Preliminary cluster 5 markers

| Gene | avg log2FC | Adjusted P | Cluster-5 detection | Other clusters |
|---|---|---|---|---|
| Cck | 4.78 | $1.09 \times 10^{-153}$ | 20.8% | 1.3% |
| Mmp10 | 4.26 | $2.90 \times 10^{-186}$ | 34.4% | 3.8% |
| Spp1 | 3.97 | $2.60 \times 10^{-214}$ | 72.1% | 20.7% |
| Cpne4 | 3.72 | $4.72 \times 10^{-81}$ | 12.8% | 1.0% |
| Gjb3 | 3.54 | $8.60 \times 10^{-93}$ | 14.6% | 1.2% |
| Mmp3 | 3.33 | $1.46 \times 10^{-130}$ | 56.8% | 18.0% |
| Etv4 | 3.28 | $2.08 \times 10^{-73}$ | 12.3% | 1.1% |
| Ctxnd1 | 3.02 | $8.88 \times 10^{-89}$ | 18.7% | 2.3% |
| Mmp9 | 2.99 | $2.61 \times 10^{-159}$ | 58.9% | 15.9% |
| Ass1 | 2.92 | $2.10 \times 10^{-234}$ | 56.0% | 9.3% |

#### SI Fig 3. PDGFRA is retained across fibroblast and early fate-associated states in large wounds

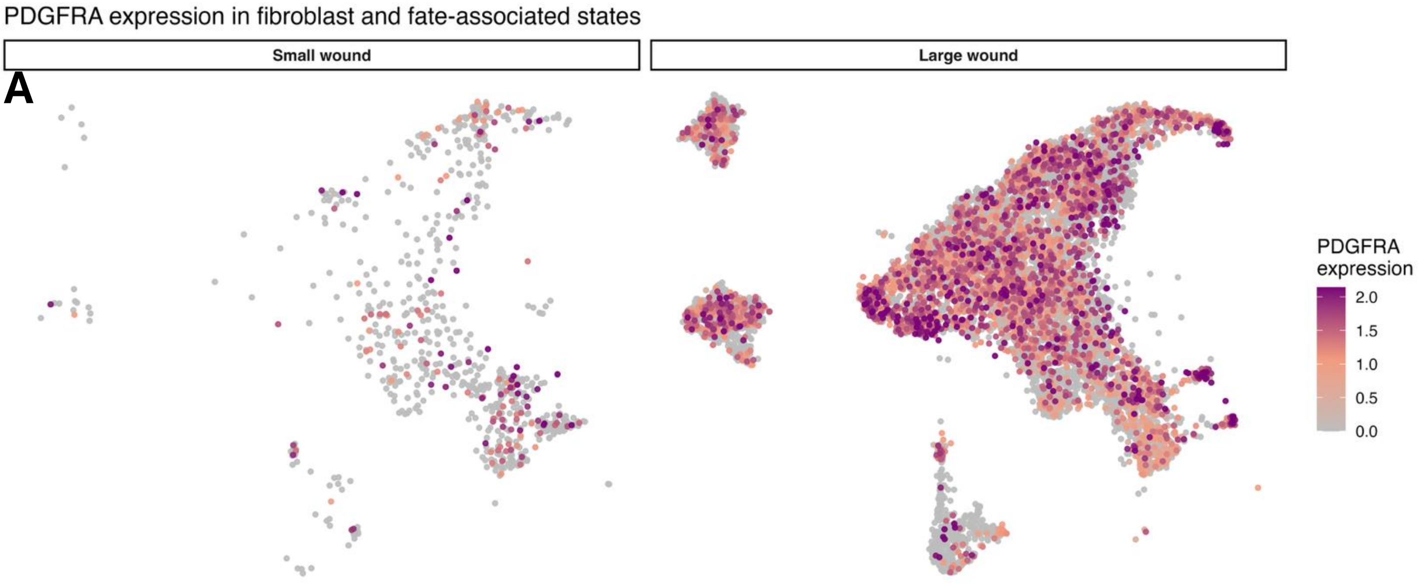


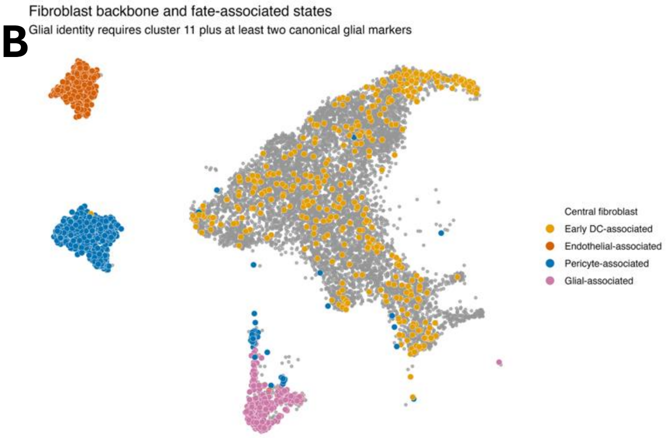

**SI Fig 4. NCAM1 marks competence-associated fibroblast states enriched during large-wound regeneration**

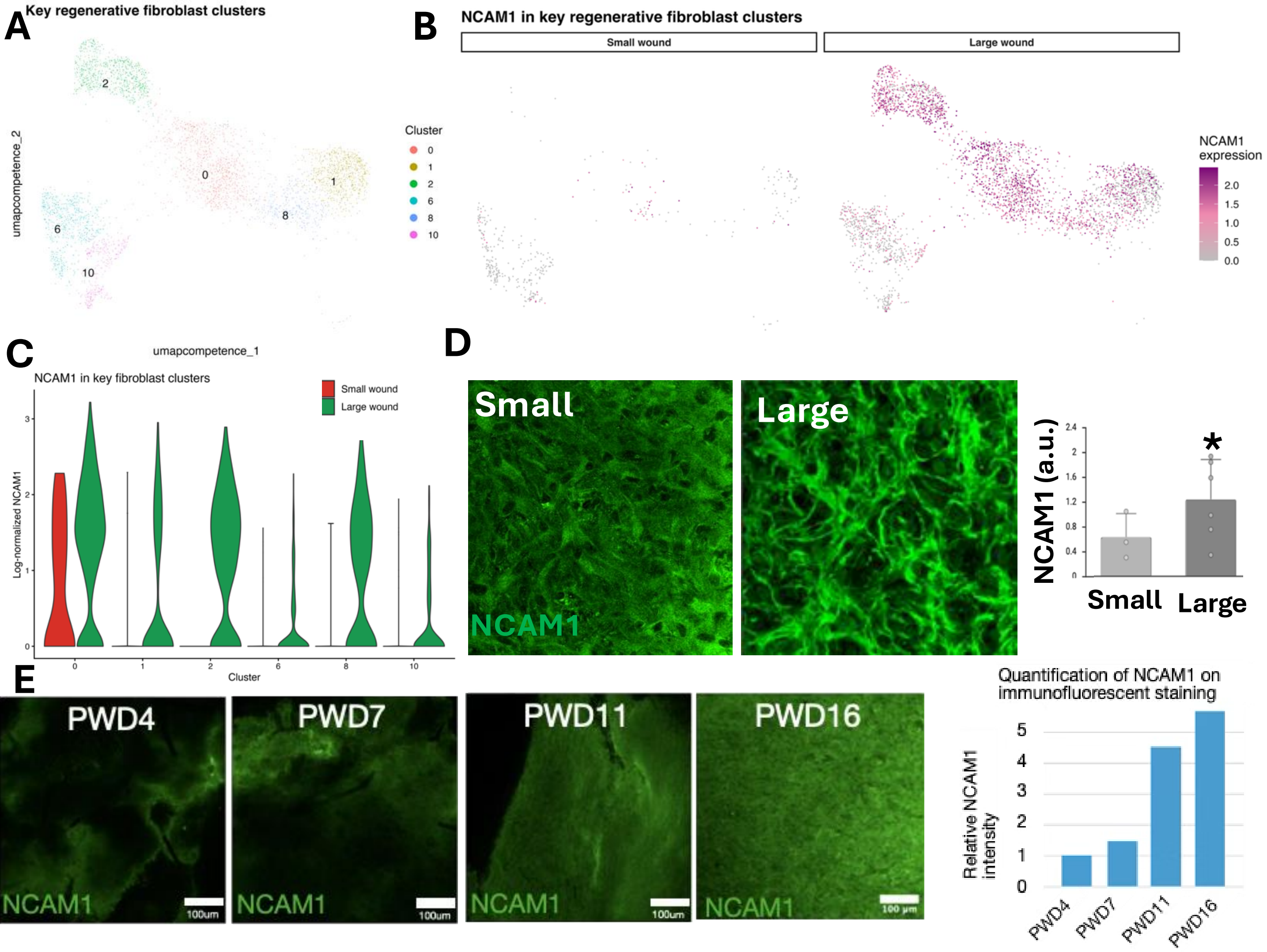

**SI Fig 5. C2 fibroblasts exhibit a transient signal-provider and matrix-remodeling program**

A

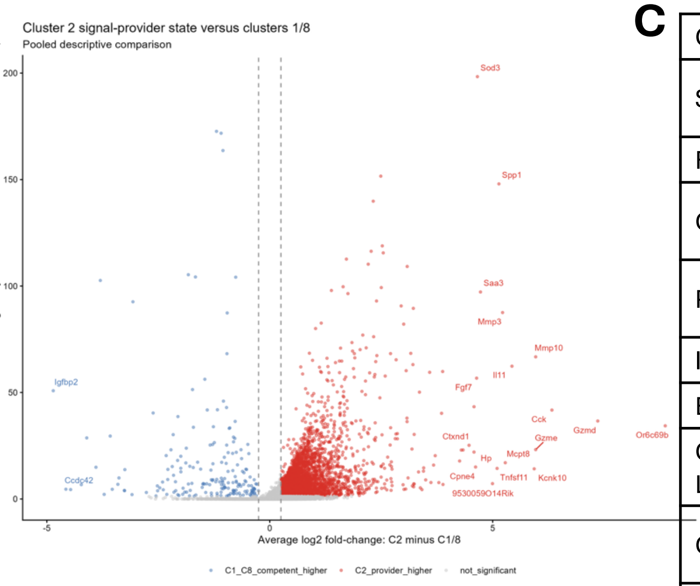


B

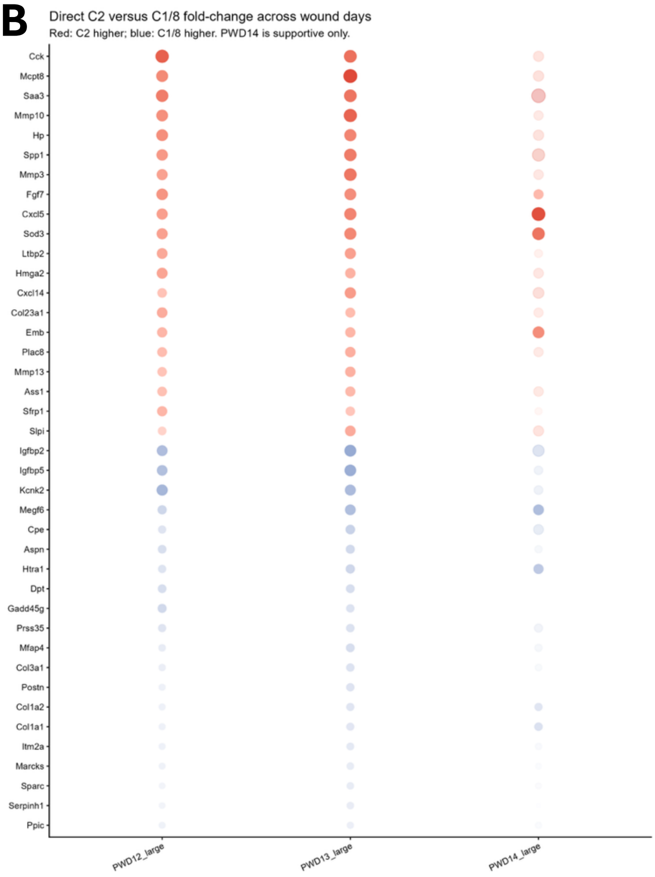


C

| Candidate | Potential function |
|---|---|
| SPP1 | Integrin/CD44 signaling, adhesion, migration and matrix remodeling |
| FGF7 | Paracrine growth and survival signaling |
| CXCL12 | Recruitment, positioning and maintenance of responsive cells |
| PDGFA | PDGFRA-positive fibroblast survival/activation |
| INHBA | Activin signaling and state regulation |
| EREG | EGFR-mediated growth and plasticity |
| CXCL5/CXCL16 | Chemotactic and inflammatory niche signaling |
| CCL7/CCL8 | Recruitment and inflammatory communication |
| THBS1 | Matrix-integrin/CD47-mediated signaling |
| TNC | Matrix-mediated signaling and plasticity |
| GREM1 | BMP antagonism and alteration of morphogen availability |
| SERPINE2 | Extracellular protease regulation |
| MMP3/9/10 | Matrix remodeling and release of matrix-bound signals |

D

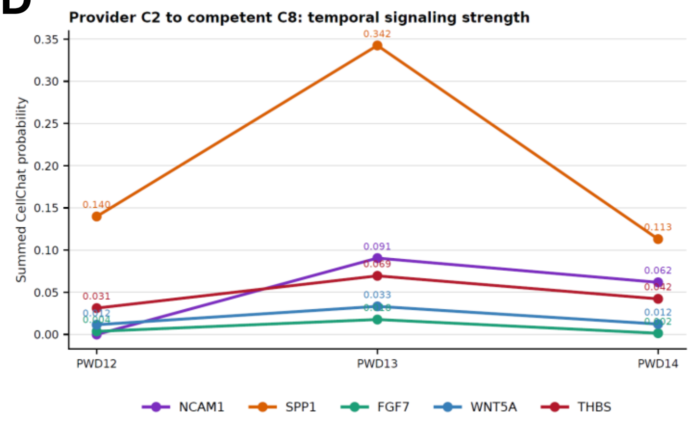

**SI Fig 6. Fibroblast-centered trajectories show progressive acquisition of four regenerative fate programs**

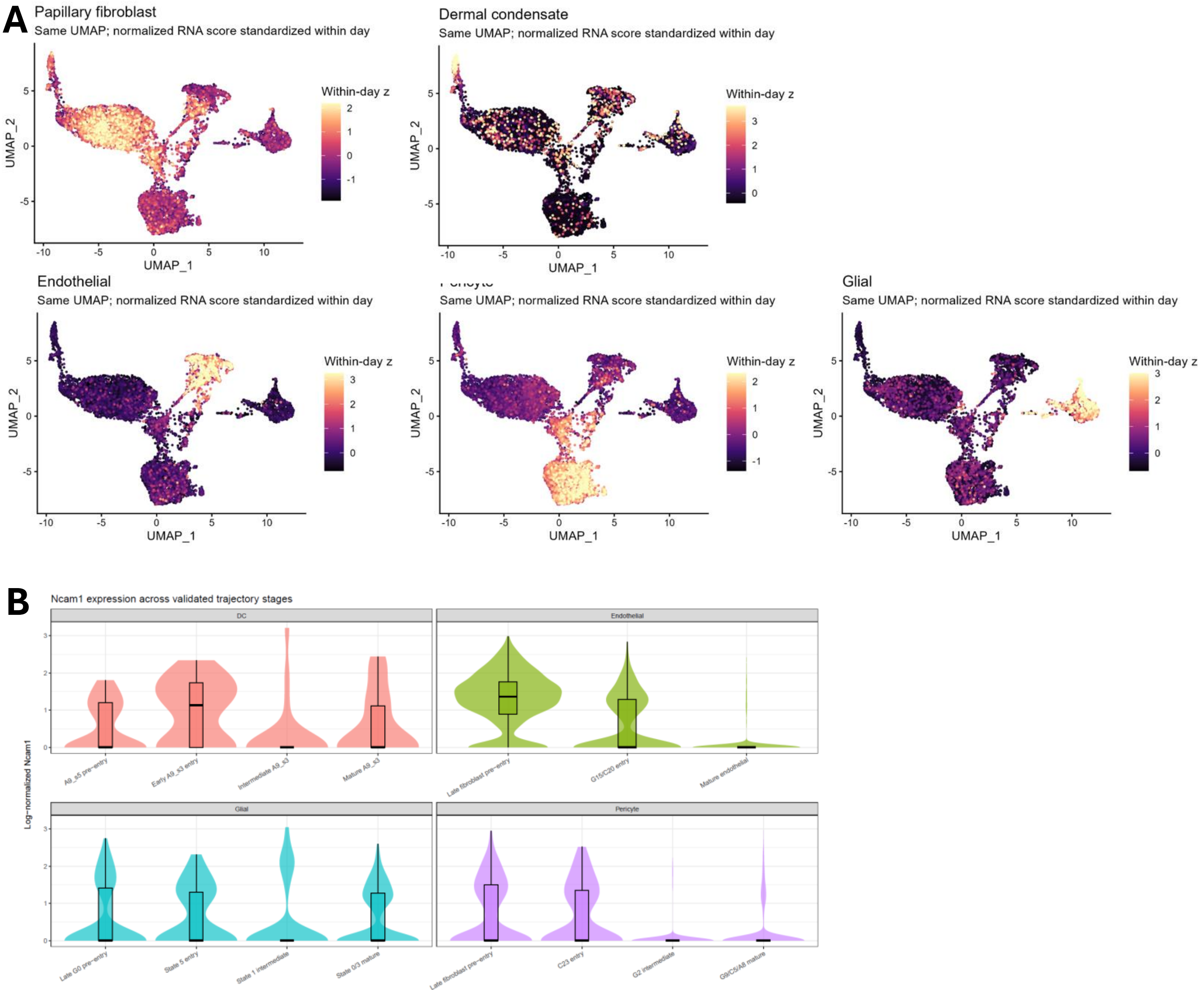

**SI Fig 7. C2 signaling fibroblasts provide shared and branch-associated inputs to competent and fate-entry states**

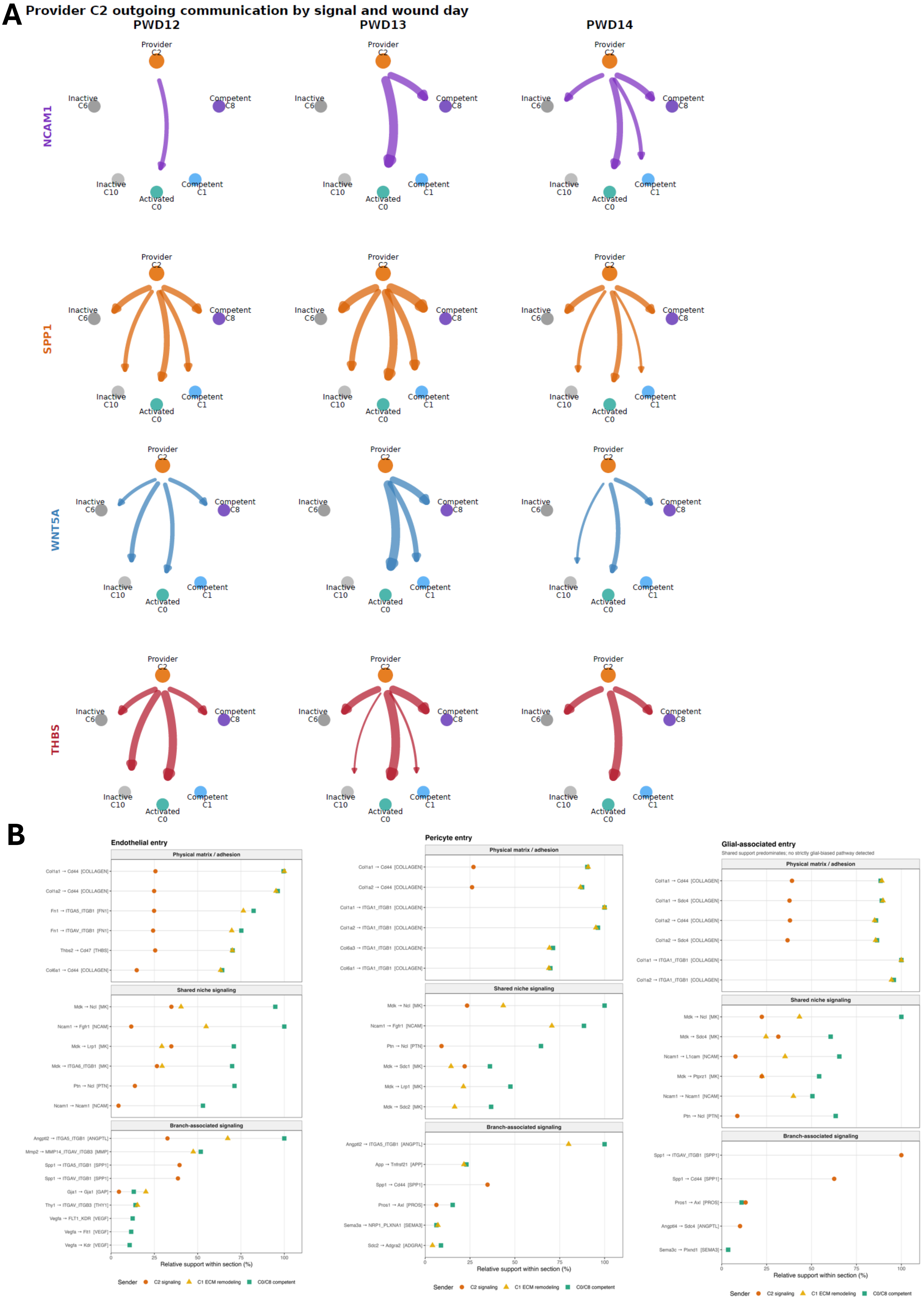

#### SI Fig 8. Gene modules define staged loss of fibroblast identity and acquisition of regenerative fates

**A**

| Module / readout | Branch | Genes | Gene count | Purpose |
|---|---|---|---|---|
| Fibroblast identity | Shared | Pdgfra, Col1a1, Col1a2, Col3a1, Dcn, Lum, Col6a1, Col6a2 | 8 | Tracks retention or loss of fibroblast identity across every branch. |
| Dermal condensate / DP acquisition | Dermal condensate | Lef1, Sox2, Fgf10, Bmp4, Rspo1, Wif1, Corin, Alpl | 8 | Tracks acquisition of regenerative dermal-condensate/dermal-papilla identity. |
| Endothelial acquisition | Endothelial | Kdr, Klf2, Esam, Emcn, Cdh5, Pecam1, Erg, Egfl7 | 8 | Tracks endothelial fate acquisition and maturation. |
| Pericyte acquisition | Pericyte | Rgs5, Cspg4, Mcam, Pdgfrb, Notch3, Des, Acta2, Cnn1 | 8 | Tracks mural/pericyte fate acquisition and maturation. |
| Glial-associated acquisition | Glial-associated | Ngfr, Fabp7, S100b, Sox10, Plp1, Mpz, Mbp, Uchl1 | 8 | Tracks glial-associated fate acquisition and maturation. |
| NCAM1 | Cross-branch single-gene readout | Ncam1 | 1 | Analyzed separately on one shared expression scale; it is not averaged into the fate modules. |

**B**

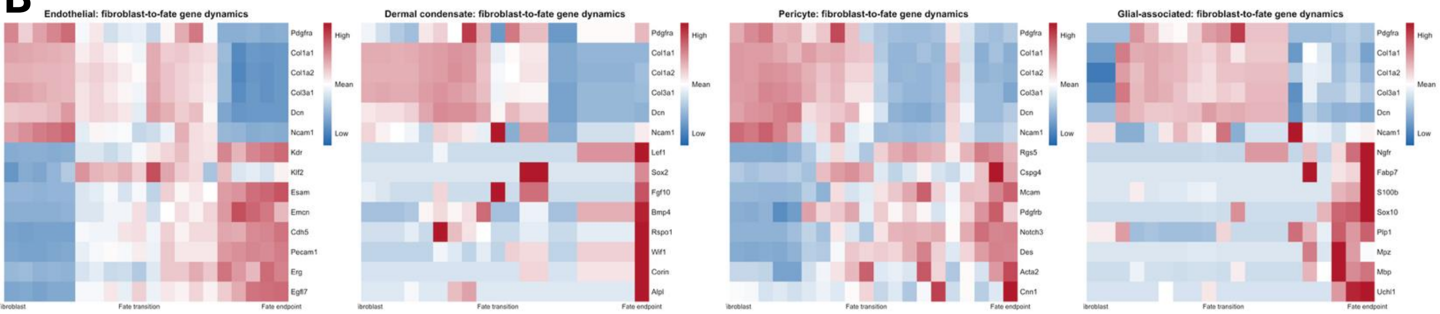


**C**

| Trajectory | Detailed states | Main-figure state |
|---|---|---|
| DC/DP | G8 fibroblast; A9_s5 branching fibroblast | Fibroblast/source |
| DC/DP | Early A9_s3; intermediate A9_s3 | Fate transition |
| DC/DP | Mature regenerative A9_s3 | Mature fate |
| Endothelial | C1-proximal fibroblast | Fibroblast/source |
| Endothelial | G15 entry; C20 intermediate | Fate transition |
| Endothelial | C9/A15 endothelial | Mature fate |
| Pericyte | Branch-proximal competent fibroblast | Fibroblast/source |
| Pericyte | C23 entry; G2 intermediate | Fate transition |
| Pericyte | G9/C5/A8 pericyte | Mature fate |
| Glial-associated | G0 glial-biased fibroblast | Fibroblast/source |
| Glial-associated | State 5 entry; state 1 intermediate | Fate transition |
| Glial-associated | State 0 repair; state 3 myelinating | Mature fate |

**D**

| Trajectory | Fibroblast/source state | Fate-transition module | Mature-fate module |
|---|---|---|---|
| DC/DP | Shared fibroblast module; includes G8 | Lef1, Fgf10, Bmp4, Rspo1 | Sox2, Wif1, Corin, Alpl |
| Endothelial | Shared fibroblast module; C1-proximal cells remain fibroblasts | Kdr, Esam, Emcn, Egfl7 | Cdh5, Pecam1, Erg, Klf2 |
| Pericyte | Shared fibroblast module; competent C1/C8-proximal cells | Rgs5, Cspg4, Mcam, Pdgfrb | Notch3, Des, Acta2, Cnn1 |
| Glial-associated | Shared fibroblast module; G0 remains fibroblast-associated | Ngfr, Fabp7, S100b, Uchl1 | Sox10, Plp1, Mpz, Mbp |

**SI Table 1. Provisional identities and condition composition of integrated small- and large-wound cell clusters**

| Cluster | Provisional identity | Small wound | Large wound |
|---|---|---|---|
| 0 | Dermal fibroblasts; mixed proliferative/contractile states | 2.52% | 24.19% |
| 1 | Dermal fibroblasts | 6.72% | 17.70% |
| 2 | Proliferating cells | 17.94% | 1.26% |
| 3 | Endothelial cells | 3.05% | 12.02% |
| 4 | Suprabasal/activated keratinocytes | 11.60% | 0.25% |
| 5 | Inflammatory macrophages | 1.24% | 9.48% |
| 6 | Antigen-presenting cells/macrophages | 6.60% | 4.11% |
| 7 | Hair-follicle/keratinocyte population | 10.11% | 0.01% |
| 8 | Antigen-presenting cells | 0.14% | 9.56% |
| 9 | Suprabasal/activated keratinocytes with some fibroblasts | 2.40% | 7.08% |
| 10 | Basal keratinocytes | 7.93% | 0.11% |
| 11 | T/NK cells | 2.45% | 4.90% |
| 12 | Antigen-presenting/dendritic cells | 2.03% | 3.82% |
| 13 | Spinous-layer keratinocytes | 5.32% | 0.40% |
| 14 | Basal keratinocytes | 5.00% | 0.18% |
| 15 | Mixed fibroblast/inflammatory population | 3.02% | 0.94% |
| 16 | Neutrophils | 2.98% | 0.27% |
| 17 | M2-like macrophages | 2.90% | 0.18% |
| 18 | Small-wound immune/epithelial population | 2.81% | 0.00% |
| 19 | Proliferating cells with fibroblast contribution | 0.01% | 2.58% |
| 20 | Minor suprabasal/pDC-associated population | 1.65% | 0.11% |
| 21 | Minor inflammatory population | 1.58% | 0.01% |
| 22 | Fibroblast-associated transitional population | 0.00% | 0.83% |

### SI Table 2. Integrated small and large wound fibroblast-only cell cluster identity

**Fibroblast competence landscape: cluster identities and dataset composition**

*Percentages are the fraction of all retained fibroblast-state cells within each dataset. Identities are based on markers, prior-state crosswalks, and post hoc module scores.*

| Cluster | Proposed identity | Role in competence model | Confidence | Key markers | Interpretive evidence | Total cells | Small combined | PWD12 large | PWD13 large | PWD14 large |
|---|---|---|---|---|---|---|---|---|---|---|
| 0 | NCAM1/PTN activated fibroblast | Activation-to-competence intermediate | Moderate | Pim1, Fabp5, Ncam1, Ptn, Igfbp7, Lrrc15, Cxcl2 | Mixed G0/G11/G1/C1; activated but less specifically tied to the core competence sources than clusters 1/8. | 996 | 8.23% | 22.34% | 16.04% | 5.21% |
| **1** | **HTRA1/CRABP1 early competent fibroblast** | **Core early competence candidate** | **High** | **Crabp1, Htra1, Igfbp5, Megf6, Prss35, Col7a1, Mfge8** | **Major shared state of PWD12 G8 and PWD13 C1; retains strong fibroblast identity.** | **741** | **7.49%** | **17.57%** | **12.00%** | **2.17%** |
| **2** | **L5/C13 deep competence signaling hub** | **Deep competence / signaling-remodeling hub** | **High** | **Ptprn, Sod3, Pla2g7, Tnc, Mmp10, Spp1, Il11, Ccn4, Hmga2, Inhba** | **Reconstructs PWD13 C13/L5 and a smaller PWD12 G0 population; nearly absent from Small.** | **737** | **0.90%** | **8.58%** | **26.62%** | **0.93%** |
| 3 | PTN/MDK trophic fibroblast | Trophic ECM state; competence-adjacent | Moderate | Ptn, Mest, H19, Mdk, Igfbp4, Gpx3, Cxcl1 | Dominated by PWD12 G1; not a primary C1/G0/G8/C13 competence cluster. | 707 | 8.83% | 21.59% | 1.15% | 5.68% |
| 4 | ASPN/CDH3 matrix-organizing fibroblast | PWD14-enriched remodeling state | Moderate | Aspn, Cdh3, Omd, Lgr5, Serpinh1, Cdo1, Lrrc17 | Predominantly PWD14 A4 with a smaller PWD13 C1 contribution; not selected as the core early-competence reference. | 657 | 1.05% | 3.05% | 8.54% | 21.37% |
| 5 | IGFBP2/CYP26B1 G8/S2-related fibroblast | Competence-adjacent retinoid-responsive state | Moderate-high | Igfbp2, Ppp1r14a, Cyp26b1, Fgfr4, Nrn1, Ntn1, Lgals3 | Contains PWD12 G8, Small S2, PWD13 C1 and PWD14 A4; demonstrates a related state in Small wound. | 651 | 13.92% | 8.87% | 5.92% | 12.03% |
| 6 | PI16/CLEC3B inactive root-like fibroblast | Primary inactive/root reference | High | Clec3b, Wnt2, Sfrp4, Pi16, Tnxb, Tppp3, Cd34, Scara5 | Most abundant Small state; broad quiescent fibroblast program and minimal competence-source contribution. | 573 | 24.85% | 3.85% | 3.41% | 12.91% |
| 7 | DLK1/NOV adventitial ECM fibroblast | Inactive-like/adventitial state | Moderate | Kcnj8, Nov, Egfl6, Mgp, Dlk1, Gas6, Nrk, Cpxm2, Eln | Mostly PWD14/PWD12 A0/G1-like cells; retained as a distinct fibroblast state, not a core competence cluster. | 362 | 6.59% | 3.18% | 0.52% | 11.98% |
| **8** | **TWIST/POSTN advanced competent fibroblast** | **Advanced matrix-remodeling competence** | **High** | **Crabp1, Htra1, Apcdd1, Robo2, Postn, Twist1, Twist2, Mmp11** | **Strongly derived from C1/G8/G0 and markedly depleted in Small wound.** | **343** | **1.05%** | **5.61%** | **9.33%** | **1.24%** |
| 9 | POSTN/CCN4/INHBA activated fibroblast | PWD13 C3-like activated remodeling state | Moderate-high | Postn, Ccn4, Inhba, Col6a3, Col5a1, Cemip2 | PWD13-enriched C3/C1 state with some contractile/remodeling signal; not used to define initial competence. | 335 | 0.90% | 0.50% | 15.51% | 1.08% |
| 10 | CLEC3B/INMT reticular quiescent fibroblast | Secondary inactive/root reference | High | Clec3b, Abca8a, Cygb, Inmt, Tnxb, Pi16, Tmeff2 | Quiescent/reticular state enriched in Small and PWD14 and nearly absent at PWD13. | 313 | 15.12% | 0.92% | 0.63% | 9.19% |
| 11 | SCX/TNMD tendon-like ECM fibroblast | Specialized ECM fibroblast; outside core competence | High | Fmod, Col11a1, Acan, Scx, Thbs4, Tnmd, Cilp, Matn4 | Distinct tendon/ligament-like matrix program; not considered part of the principal competence transition. | 256 | 1.95% | 3.93% | 0.21% | 7.49% |
| 12 | ALDH3A1/CCL19 specialized fibroblast | Specialized Small/PWD14 state | Moderate | Aldh3a1, Ccl19, Col15a1, Scara5, Rspo1, Bmp4, Clec3b | Mostly PWD14 A9_s5 and Small S9; excluded from the initial inactive-versus-competent contrast. | 183 | 8.68% | 0.00% | 0.10% | 6.35% |
| 13 | SMPD3/PGR rare specialized fibroblast | Rare PWD14-restricted state | Low-moderate | Smpd3, Pgr, Cxcl13, Krt19, Aldh1a3, Tek, Ackr1 | Small cluster dominated by PWD14 A9_s6; possible vascular association/contamination requires caution. | 49 | 0.45% | 0.00% | 0.00% | 2.37% |
